# Differential retinoic acid responses across testicular development *in vitro*

**DOI:** 10.64898/2025.12.30.696417

**Authors:** Brad C. Hansen, Sarah A. Hatem, Lucas A. Huang, John K. Amory, Elaine M. Faustman, Edward J. Kelly

## Abstract

Spermatogonial differentiation is controlled by distinct spatial and temporal activities of metabolizing enzymes and signaling molecule formation. In vitro models are reductionist ways to examine the interactions between Sertoli cells and spermatogonial stem cells, which may be therapeutic targets for infertility that cause non-obstructive azoospermia, due to either Sertoli-cell only syndrome, hypospermatogenesis or maturation arrest. Here, we find that nanomolar doses of isotretinoin are sufficient to drive *Stra8* expression in vitro, an interaction that is both dose-dependent and inhibitable. We compare complex in vitro models (CIVMs) seeded from cells isolated post-natal day 5 (PND 5) and post-natal day 10 (PND 10) Sprague Dawley rat testis. The CIVMs maintain metabolic capacity to produce bioactive retinoids form retinol. We also investigate the impact of common media supplementations on spermatogonial phenotype and find that they can impact the expression of *Stra8* and *Plzf* as markers of early differentiating and undifferentiated spermatogonia, respectively. These results highlight the power of *in vitro* models to investigate the dynamics of the spermatogonial niche.

## Introduction

The testis remains an elusive target for new complex in vitro models (CIVMs), despite 9% of males experiencing infertility in their lifetime globally and difficulty in identifying testicular impacts during toxicology screening and drug development [1–4]. A considerable proportion (∼40%) of infertility cases are idiopathic [5]. These cases require increased research in the field and particular emphasis on critical developmental windows during testicular development. Neonatal development of testis (post-natal days 3-7 in the rat) is critical for the establishment of the testis niche [6], where dysfunction of neonatal testicular development is common with lasting adverse effects on reproductive health and fertility [7–10].

*In vitro* models of the testis need to consider the complexity of the organ where there are two distinct functional areas: the seminiferous tubule, the site of spermatogenesis, and the interstitial space, the site of testosterone biosynthesis and interaction with vasculature. Sertoli cells are the supporting somatic cells of the seminiferous tubule, responsible for maintaining spermatogenesis through close contact with spermatogonia and establishment of the immune privilege area for post-meiotic cells [11]. Sertoli cells continue to proliferate early in life, ceasing around post-natal day (PND) 15 in rats [12,13]. Spermatogonia reside at the basal lamina of the seminiferous tubule in close contact with the Sertoli cells, where they undergo signaling towards either proliferation (renewal) or differentiation and commitment to meiosis. Retinoic acid (RA) is the well-established initiator of spermatogonial differentiation and most bioactive RA in testes (all-trans RA; *at*-RA) is produced within the testis, where ALDH1A enzymes are present within the Sertoli cells of the neonatal testis to form bioactive RA and CYP26 enzymes are present within the peritubular myoid cells that encircle the tubules to form a barrier to exogenous retinoic acid and the pre-mature initiation of spermatogenesis [14].

Mechanistic studies on the interactions between Sertoli cells and spermatogonial stem cells are critically needed to study idiopathic infertility conditions that cause non-obstructive azoospermia, due to either Sertoli-cell only syndrome (SCOS), hypospermatogenesis, or maturation arrest. Both SCOS and maturation arrest are significant sources of clinical infertility from non-obstructive azoospermia where there is either an arrest of the spermatogonial cells in their pre-meiotic phenotype or an absence of spermatogonial cells altogether (SCOS). In both SCOS and maturation arrest, genetic and environmental factors are implicated, supporting the use of *in vitro* models where gene x environment interactions are controllable [15–17].

In this study, we compare CIVMs derived from post-natal day 5 (PND 5) and post-natal day 10 (PND 10) Sprague Dawley rat testis. At PND 5, these are neonatal rat testis, where Sertoli cells are actively proliferating and gonocytes are transitioning into type A undifferentiated (type A) spermatogonia [6]. By PND 10, these are now early infantile rat testis where both Sertoli cells and spermatogonia are actively proliferating, alongside the emergence of early differentiating (type B spermatogonia) [6]. While both PND 5 and PND 10 Sprague Dawley rats are reproductively immature animals, the likely emergence of type B in the PND 10 testis and the impact of this difference in spermatogonial phenotype, is of particular interest in this study.

## Methods

### Tissue isolation and culture establishment

Testes were isolated from neonatal Sprague Dawley rats (Strain 001, Charles River) on post-natal day 5 (PND 5) or post-natal day 10 (PND 10) using sterile forceps and placed in Minimum Essential Media (MEM, Gibco) containing 1% penicillin-streptomycin (PS, Gibco). Testes in MEM with 1% PS were kept on ice (< 30 minutes) and moved to a sterile Biological Safety Cabinet (BSC) for the remaining tissue isolation. When moved into the BSC, the testes were washed with fresh MEM twice by inversion in a conical tube. The washed testes were then placed in a 6-well cell culture plate with 2-3 mLs of cold MEM in each well. Using sterile forceps, the testes were placed in a well, the plate swirled, and the testis moved to the next well, as a wash. After two washes, the plate was moved under a dissecting microscope in the BSC, where the epididymides were removed and each testis detunicated. Each detunicated testis was fragmented into 3-4 pieces and moved to a well with fresh MEM. Then, 2-3 fragments were moved to cryovials with 1mL freezing medium (70 mmol/L sucrose diluted in phenol-red free DMEM/F12 with 10% v/v DMSO; Gibco; freezing media recipe from [18]). The vials were moved to a controlled freezing device (Mr. Frosty) and kept in a −80C freezing for 24 hours, after which the cryovials were moved to liquid nitrogen for long term storage.

For cell plating, the vials were removed from liquid nitrogen storage and rapidly thawed in a water bath (37C). The fragments were removed from the cryovials into a conical tube with fresh MEM using a transfer pipette to minimize freezing media crossover. The fragments were allowed to settle on ice for 10 minutes before sequential enzyme digestion using hyaluronidase and collagenase; additional enzymatic digestion protocol details in published protocol [19]. The digested tissue was further dissociated with a brief digestion in 0.05% Trypsin-EDTA solution (Gibco), followed by centrifugation to resuspend in complete testis cell culture medium (containing phenol-red-free MEM [Gibco], 1% sodium pyruvate [Gibco], 1% non-essential amino acids [Gibco], 1% ITS+ culture supplement [Corning], 1% PS [Gibco], epidermal growth factor [Sigma Aldrich], and sodium lactate [Sigma Aldrich]); complete media formulation available in published protocol [19]. The cells were resuspended at a concentration of 1.6 million cells per mL and 100uL plated in each well of a 48-well plate that had previously been coated with a 1% v/v Matrigel in MEM solution for > 1 hour at room temperature. Immediately after adding the cell suspension to each well, an additional 100uL of Matrigel overlay solution (complete testis medium with 400ug/mL Matrigel protein), was added to each well, for a final plating density of 800,000 cells per mL and 200 ug/mL Matrigel overlay in a 200 uL total culture volume. Culture media was changed 24-hours after plating, and every 48-hours thereafter.

### Retinoid dosing and inhibitor treatment

Isotretinoin (13-cis retinoic acid) and Retinol (both from Sigma Aldrich) stocks were prepared at 1 mg/mL in analytical grade ethanol and mainlined at −80C protected from light. Retinoids were diluted to working stocks in analytical grade ethanol. The working stocks were diluted in complete cell culture media to achieve the experimental doses and consistent 0.1% ethanol v/v. WIN-18446 was used as a specific inhibitor of the aldehyde dehydrogenase (ALDH) enzymes and BMS-189453 was used as a pan-retinoic acid receptor (RAR_(α,β,γ)_) antagonist. WIN-18446 (Sigma Aldrich) was diluted in DMSO to a stock concentration of 1mM. BMS-189453 (MedChem Express) was also diluted in DMSO to a stock concentration of 1mM. All control experiments contained an equal volume of the organic component as the treatment groups (Ethanol or DMSO), in the stimulus and inhibitory studies, the organic fractions were 0.1% v/v and 0.2% v/v, respectively.

### Culture media supplementation

Culture media supplementation experiments used three media supplement approaches common for *in vitro* testis models: spermatogonial stem cell (SSC) growth factors [20], lipid-rich albumin (AlbuMAX, Thermo Fisher Scientific) [21], and a defined bovine serum (Knockout Serum Replacement) [22,23]. SSC growth factor media was prepared as published [20]: using recombinant rat growth factors, 40 ng/mL GDNF (R&D 512-GF-010/CF), 1 ng/mL FGF2 (R&D 3339-FB-025/CF), and 300n/mL GFRA1 (R&D 560-GR- 100/CF) in complete testis media. Lipid rich albumin media was prepared by adding 20mg/mL of AlbuMAX I Lipid-Rich BSA (Thermo Fisher Scientific) to complete testis media and sterile filtering (0.2um filter), and defined bovine serum media was prepared by adding 10% v/v Knockout Serum Replacement (Gibco) to complete testis media. Control CIVMs were maintained in complete testis media. All cells were plated in control media and changed to treatment media 24 hours after plating, then changed every 48 hours thereafter through 15 days *in vitro*.

### RNA relative expression analyses and sequencing

Cells were lysed from the 48-well plates using 250uL RLT Lysis buffer (Qiagen) per well. The lysis buffer was added to the well, mixed thoroughly, and moved to an RNAse free microcentrifuge tube which was stored at −80C until extraction.

To perform reverse-transcription quantitative PCR (RT-qPCR), total RNA was extracted using a RNeasy Mini Kit (Qiagen), following manufacturer protocols. The quality and concentration of the extracted RNA was determined using a Nanodrop spectrophotometer (ND-1000; Thermo Fisher Scientific). Reverse transcription of the extracted RNA to cDNA was performed using an Applied Biosystems High-Capacity cDNA Reverse Transcription Kit, following manufacturer protocols and using a C1000 Touch Thermal Cycler (Bio-Rad). The reverse transcribed cDNA samples were diluted with nuclease-free water to 2.5 ng/μL and used for quantitative polymerase chain reaction using TaqMan assays and TaqMan Fast Advanced Master Mix in a QuantStudio 3 system (Applied Biosystems). Each RT-qPCR mixture consisted of 10 μL of TaqMan Fast Advanced Master Mix, 5 μL of nuclease-free water, 1 μL of TaqMan Gene Expression Assay, and 4 μL of cDNA (10ng of cDNA total per reaction). The RT-qPCR thermocycler settings followed TaqMan protocols. RT-qPCR was used to target the rat genes *Stra8* (Rn01747849_m1) and *Plzf* (Rn01418644_m1). Each sample’s gene expression was normalized using the difference in cycle threshold (ΔCt) calculation for relative quantification of gene expression to the housekeeping gene beta-actin (*Actb;* TaqMan Rn00667869_m1).

To perform bulk RNA-sequencing, the cell lysate samples were sent to Novogene America, where the RNA was isolated and messenger RNA was purified from total RNA using poly-T oligo-attached magnetic beads. The purified mRNA was fragmented and the first strand cDNA was synthesized using random hexamer primers, followed by second strand cDNA synthesis using dTTP for nondirectional library, and sequenced using the NovaSeq X Plus sequencing platform (Illumina). RNA quality was determine using a bioanalyzer, with all RNA Integrity Numbers (RINs) above 9, indicating high quality RNA for sequencing.

### Transcriptomics

Sequencing reads were aligned to the rat genome, GRCr8 (ensembl), using STAR aligner and gene counts tabulated using the STAR gene counts function (genecounts) [24]. Aligned read counts were processed using edgeR, which included removal of lowly expressed genes, normalization using the trimmed-mean of M-values method [25], and comparisons made between PND 5 and PND 10 CIVMs using a Quasi-likelihood F-test [26]. Statistically significant gene expression was determined based on a false discovery rate (FDR) less than 0.05 and a log2 fold change of at least 0.5 in either direction.

Pathway enrichment analyses were calculated using the TopGO package (v. 2.60.1; [27]). This method calculated the overrepresentation of differentially genes, as determined by edgeR analyses, annotated in pathways within the Biological Processes family of the Gene Ontology database [28,29]. This analysis considers if a pathway contains more genes that are differentially expressed than would be expected based on the rate of differentially expressed genes in the dataset, the statistical significance is calculated with a fishers exact test to identify how likely the observed differential gene proportion would be if the genes were selected at random.

Transcription factor enrichment assessment was completed using the decoupleR package (v. 2.9.7, [30]). The *run_ulm* function was used for univariate linear modeling on the matrix of differentially expressed genes where a synthetic variable (*t*), was created by multiplying the log2 fold change by the −log10(p-value) to incorporate magnitude, directionally, and statistical significance into a single metric for modeling. This follows the standard protocol with the *run_ulm* function, and a minimum of ten genes were required to be annotated for each transcription factor. The transcription factor and gene associations were downloaded from the omnipath database (rat specific database #10116; [31,32]). The omnipath database data was not specific to any developmental stage. The transcription factor activities with a p-value of less than 0.05 based on the fitted univariate linear model were considered.

### Deconvolution

Deconvolution estimates of the cell type proportions were completed using the *run-DWLS* and *run_OLS* functions within the DWLS package (v. 0.1.0) to implement the dampened weighted least squares and ordinary least squares regressions, respectively [33]. These regression-based deconvolution methods require two inputs: a library-size normalized matrix of gene counts from bulk RNA-seq samples and a signature matrix where each cell type expected to be within the bulk RNA-seq sample is represented by a gene expression values, these values are the product of a regression on a larger set of cells annotated to that cell type from publicly available single cells dataset (see supplemental table 2 for details on single-cell references used here). Four single cell references were used in these analyses: two from neonatal human samples and two from neonatal mouse samples (supplemental table 2). The cell types were annotated from the publicly available single-cell references to the following groupings: Sertoli cells, Leydig/stroma cells, germ cells, and peritubular myoid cells. The germ cell fraction was defined by general germ cell marker expression and was a combination of subtypes: gonocytes, pro-spermatogonia, spermatogonial stem cells, pro-spermatogonia, as well as possible early differentiating spermatogonia. To more accurately represent the gene distributions present in a bulk RNA-seq dataset, single-cell gene expression imputation implemented through the Adaptively-thresholded Low Rank Approximation (ALRA) algorithm was applied to the single-cell references in two of the three methods used here [34]. The three methods used in this report are: dampened weighted least squares regression with an ALRA imputed reference dataset, dampened weighted least squares regression with a non-imputed reference dataset, and ordinary least squares regression with an ALRA imputed dataset. See [35] for discussion and benchmarking of deconvolution approaches for neonatal tests bulk RNA-seq. The deconvolution estimates are presented as proportions of the total cell type composition. To measure significant differences in cell type composition between the two developmental stages, PND 5 and PND 10, was assessed with an unpaired Wilcoxon ranked sum test of the estimated proportions across methods and references.

### Fluorescent microscopy

All fluorescent microscopy images presented as indicative of representative examples using indirect immunocytochemistry (ICC) with commercially available antibodies. For the ICC of the 48-well culture plates, the plates were fixed with 4% paraformaldehyde solution (PFA) for 15 minutes before washing with D-PBS+/+ (Gibco). Plates were kept at +4C until processing using indirect antibody binding. All antibody buffers were made in D-PBS +/+. The plates were removed from the +4C fridge and warmed to RT. The cells were permeabilized for 3 minutes using 0.1% Triton X-100 (Sigma Aldrich), followed by non-specific binding blocking at RT in a solution of 1% goat serum (Vector Labs) and 0.1% Tween-20 (Sigma Aldrich). After blocking, the cells were washed with 0.1% Tween-20, followed by addition of primary antibodies at a 1:100 v/v dilution in 1% goat serum and 0.1% Tween-20. The cells were moved to +4C to incubate overnight. The cells were washed three times with 0.1% tween, then moved to secondary antibody incubation at a 1:1000 v/v dilution in 1% goat serum and 0.1% Tween-20 for one hour at RT. After incubation, the cells were washed three times more with 0.1% tween, followed by incubation with a nuclear counterstain (Hoechst 33342, Thermo Fisher Scientific) for 10 minutes. The counterstained cells were washed with fresh D-PBS +/+ and stored in the dark at +4C until imaging.

Whole mount tubule imaging was performed similarly to the ICC of the cell culture plates with major modifications (tubule protocol adapted from [36]). The whole tubules isolated from detunicated neonatal rat testis were fixed in 4% PFA for two hours at +4C. After fixation, the tubules underwent heat activated antigen retrieval in a 10 mM sodium citrate buffer with 0.05% v/v Tween-20 (adjusted to pH of 6), for 15 minutes at +95C in microcentrifuge tubes using a block heater. The tubules were permeabilized twice, each for 15 minutes. The primary antibody incubations were extended to three days at +4C and the secondary antibody incubations extended to two days at +4C. The serum concentration in antibody buffers was increased to 5% v/v and washes were extended to one hour of submersion in 0.1% Tween-20 buffer three times. Nuclear counterstain incubation was extended to 30 minutes. Tubules were mounted whole on standard microscopy slides and covered with a standard #1.5 coverslip in prolong gold mounting media (Thermo Fisher scientific).

Fluorescent microscopy of 48-well plates was completed on an inverted Nikon TI-Eclipse microscope, widefield scans were completed using a Nikon TI- High-Resolution Widefield microscope, and whole mount tubule imaging was completed using an Olympus Fluoview-1000 confocal microscope.

## Results

### Transcriptomics analyses

Comparison of the transcriptomes of CIVMs derived from PND 5 and PND 10 tissue reveal significant differences between the developmental stages. CIVMs seeded with cells from three PND 5 tissue isolations were compared against CIVMs from three PND 10 tissue isolations. Each sample was a mixture of four CIVMs, representing a single biological replicate. The expression of canonical germ cell marker genes was more variable in the PND 5 derived CIVMs compared to the PND 10 CIVMs, somatic cells markers were stable across both developmental stages (Figure 1.A). Based on a significance threshold of log2 fold change greater than 0.5 or less than −0.5 and false discovery rate (FDR) threshold of less than 0.05, 850 genes were differentially expressed between the developmental stages (268 genes upregulated in PND 5 CIVMs and 582 genes upregulated in PND 10 CIVMs; Figure 1.B). Multidimensional scaling shows a higher intra-group distance between the PND 5 samples compared to the PND 10 samples, though the developmental stages remain clustered by age (Figure 1.C). Gene Ontology (GO) over-representation analysis identified that 1,337 annotated pathways (of 6,077 total annotated pathways) were over-represented, based on a fisher statistic less than 0.05, among the differentially expressed genes. A keyword search of the enriched GO terms using, ‘retino,’ ’male,’ ‘testis’ or ‘gonad’ identified nine pathways with GO terms containing these keywords that are over-represented among the differentially expressed genes between CIVMs from PND 5 and PND 10 tissue (Table 1). Critically, these overrepresented pathways include regulation of gonad development, metabolism, and responses to retinoids, as well as responses to gonadotropins. An additional 15 GO gene sets containing the keywords, ‘sertoli’, ‘leydig’, ‘testos’, ‘steroid’, or ‘hormone’, were found to be overrepresented in the differentially expressed genes between CIVMS from PND 5 and PND10 (Supplemental Table 2). The enrichment of transcription factor-target genes between the two developmental stages reveals distinct transcriptional regulatory activities, with target gene upregulation in PND 5 CIVMs (in red and with positive activity score) or upregulation in PND 10 (in blue and with positive activity score) (Figure 1.D). The top 20 transcription factors in each direction, based on activity score from univariate linear modeling of the synthetic *t* statistic, are presented (Figure 1.D), the full list of transcription factors with a statically significant association to genes differentially regulated at either timepoint is included (supplemental figure 5).

**Figure 1.**
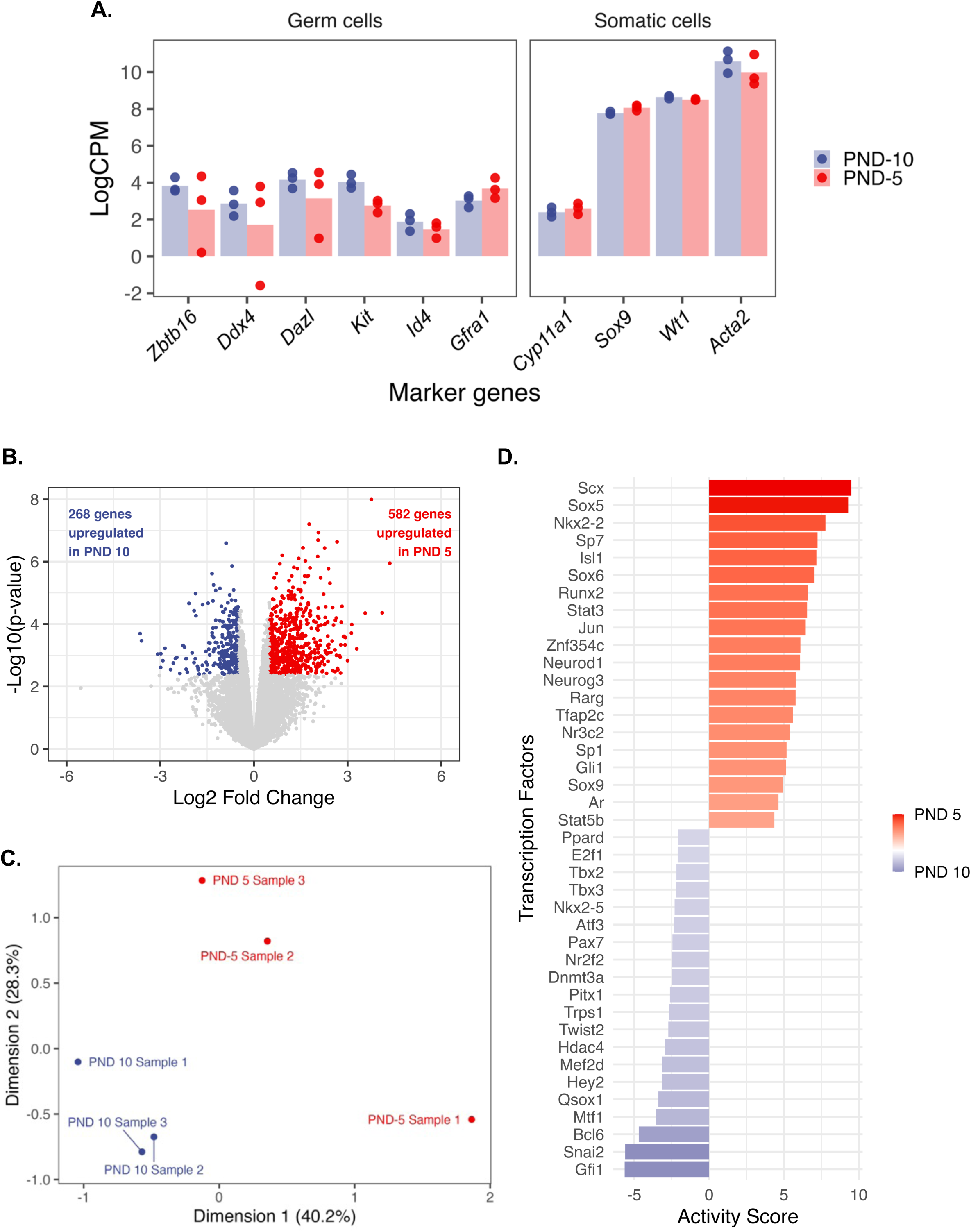
Transcriptomic Comparison of CIVMs derived from PND 5 and PND 10 after three days *in vitro*. Bulk-RNA transcriptomes were collected from PND 5 and PND 10 testis CIVMs after three days in culture. (A) The expression of canonical marker genes for germ cells in the neonatal testis alongside markers of somatic cell types. (B) Differentially expressed genes based on log2 fold change greater than 0.5 or less than −0.5 and false discovery rate (FDR) threshold of less than 0.05 shown in volcano plot with genes upregulated in PND 10 shown as a negative log2 fold change and genes upregulated in PND5 shown as a positive log2 fold change. (C) Multidimension scaling supports that the intra-developmental stage samples cluster together and suggests that the intra-sample variance is higher in PND 5 CIVMs. Percentage for each dimension represents the variance captured by that component. (D) Inferred transcription factor (TF) activity from differentially expressed genes, top 20 TFs for each developmental stage included (PND 5 in red with positive activity score, PND 10 in blue with negative activity score).

**Table 1.**
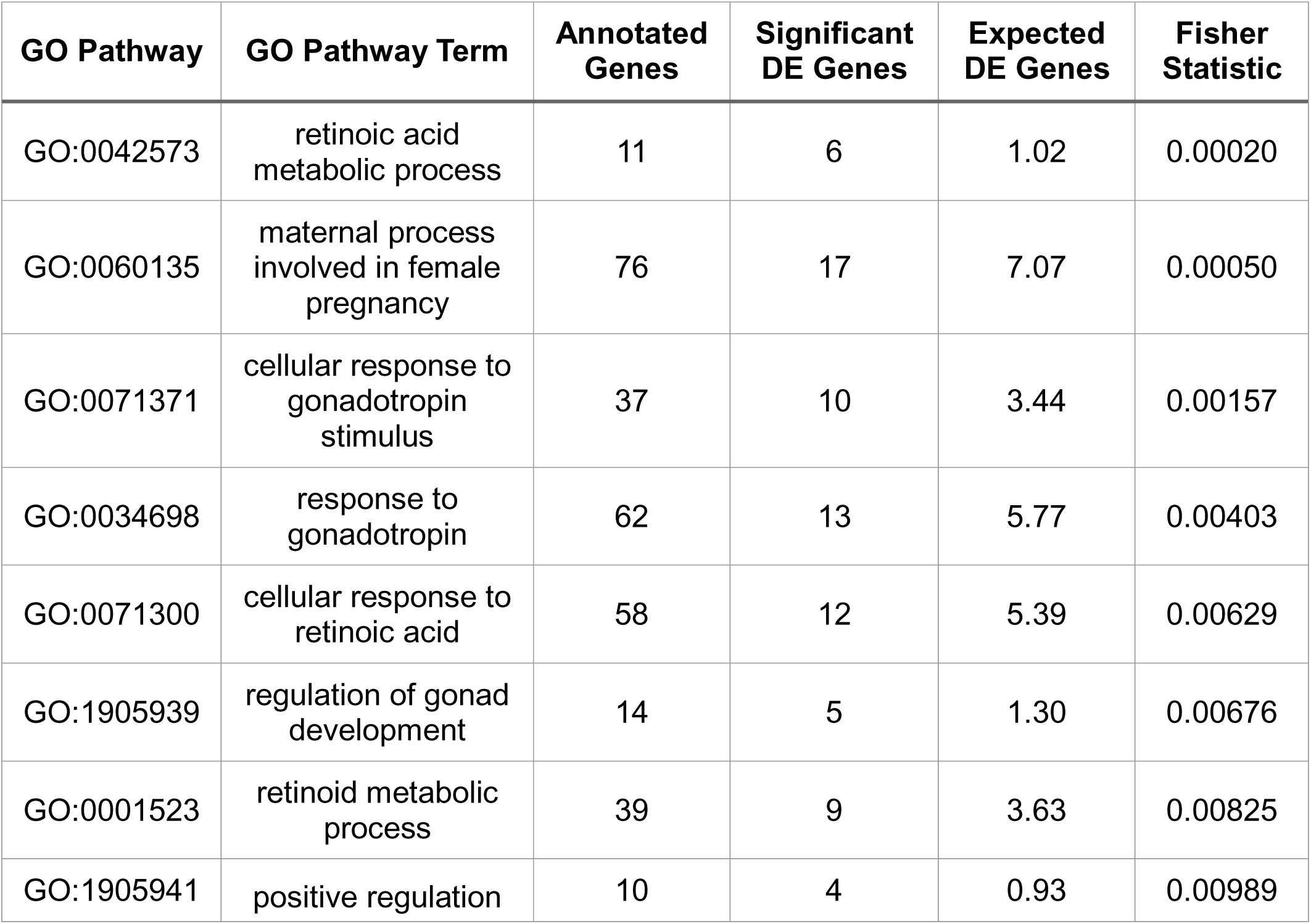
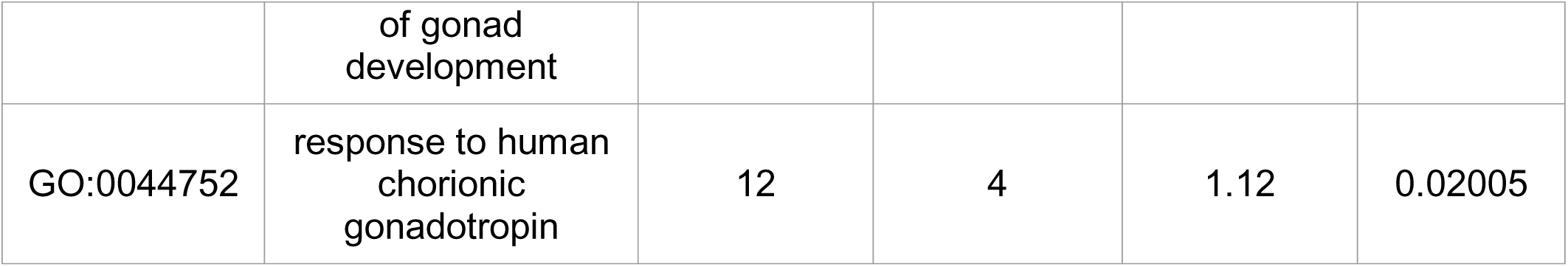
Gene Ontology overrepresentation analyses of differentially expressed genes between CIVMs derived from PND 5 and PND 10 testis.

| GO Pathway | GO Pathway Term | Annotated Genes | Significant DE Genes | Expected DE Genes | Fisher Statistic |
| --- | --- | --- | --- | --- | --- |
| GO:0042573 | retinoic acid metabolic process | 11 | 6 | 1.02 | 0.00020 |
| GO:0060135 | maternal process involved in female pregnancy | 76 | 17 | 7.07 | 0.00050 |
| GO:0071371 | cellular response to gonadotropin stimulus | 37 | 10 | 3.44 | 0.00157 |
| GO:0034698 | response to gonadotropin | 62 | 13 | 5.77 | 0.00403 |
| GO:0071300 | cellular response to retinoic acid | 58 | 12 | 5.39 | 0.00629 |
| GO:1905939 | regulation of gonad development | 14 | 5 | 1.30 | 0.00676 |
| GO:0001523 | retinoid metabolic process | 39 | 9 | 3.63 | 0.00825 |
| GO:1905941 | positive regulation of gonad development | 10 | 4 | 0.93 | 0.00989 |
| GO:0044752 | response to human chorionic gonadotropin | 12 | 4 | 1.12 | 0.02005 |

### Deconvolution analyses

The composition of CIVMs derived from each developmental stage was estimated using bulk RNA-seq deconvolution methods. Three deconvolution approaches were used, each a combination of an imputation algorithm and a deconvolution regression: DWLS deconvolution with ALRA imputed reference data, OLS deconvolution with ALRA imputed reference data, and DWLS deconvolution with non-imputed reference data. These methods were identified in previous analysis as the best for neonatal testis models (method benchmarked in [35]). Deconvolution estimates for each cell type were relatively consistent between developmental stages and no significant differences were identified through an unpaired Wilcoxon rank-sum test comparing the cell types at each developmental stage (Figure 2.A). The average estimates for cell proportion across methods and single-cell references were: 0.091 and 0.083 for germ cells, 0.116 and 0.097 for Leydig/Stromal cells, 0.309 and 0.330 for peritubular myoid cells, and 0.484 and 0.490 for Sertoli cells, in the PND 5 and PND 10 CIVMs respectively (Table 2).

**Figure 2.**
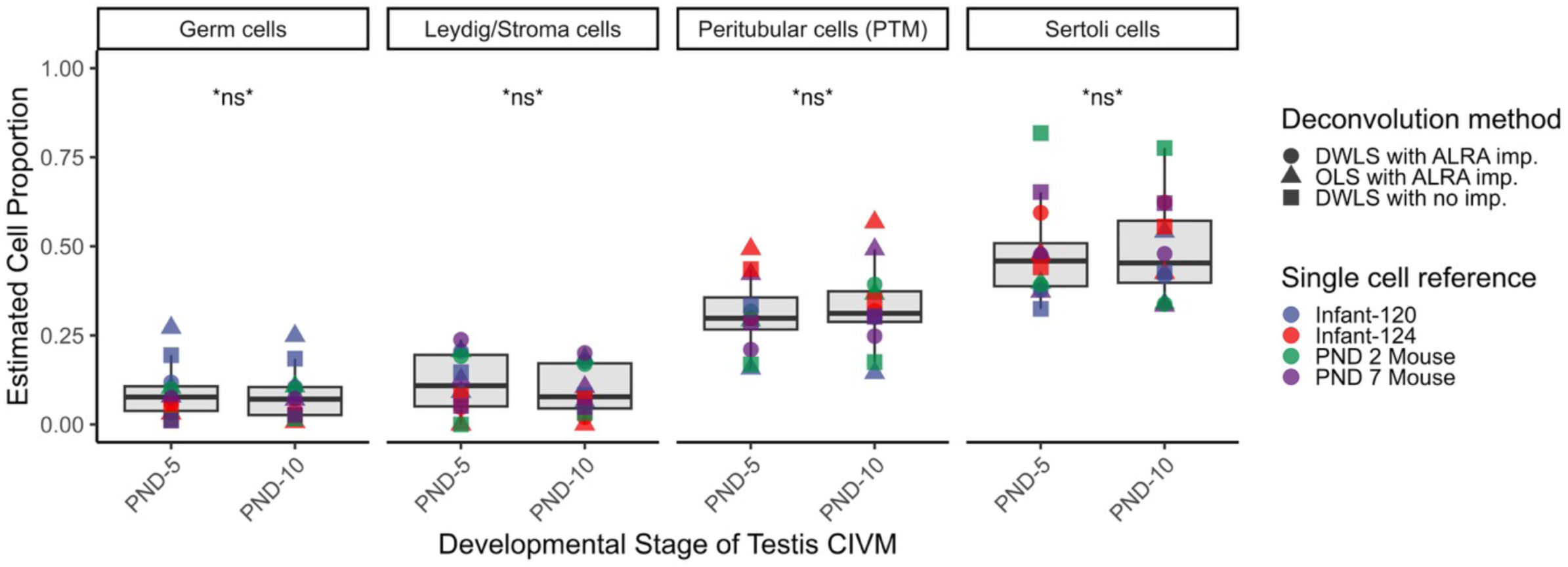
RNA-seq Deconvolution based- estimates of cell type proportion within CIVMs of neonatal testis from PND 5 and PND 10 tissue. Bulk-transcriptomes from the PND 5 and PND 10 derived CIVMs were deconvolved using publicly available methods. Shape indicated the combination of deconvolution method and color indicates the single-cell reference used. Each point is the mean of the three biological replicate CIVM RNA-seq samples for each developmental stage per each combination of method and single cell reference. Boxplots represent the full distribution of estimates (including biological replicate CIVMs). There were no statistically significant differences identified between the cell type estimates at the two stages, as determined by an un-paired Wilcoxon ranked sum test p-value greater than 0.05 for all cell types (denoted by ‘ns’).

**Table 2.** Means and ranges of RNA-seq deconvolution based- estimates of cell type proportion within CIVMs of neonatal testis from PND 5 and PND 10 tissue.

| **CIVM Developmental Stage** | **Cell Type** | **Mean Proportion Estimate** | **Range of Proportion Estimates** |
| --- | --- | --- | --- |
| PND 5 | Germ cells | 0.091 | 0.011 – 0.271 |
|  | Leydig/Stroma cells | 0.116 | 0 – 0.238 |
|  | Peritubular cells | 0.309 | 0.157 – 0.493 |
|  | Sertoli cells | 0.484 | 0.324 – 0.818 |
| PND 10 | Germ cells | 0.083 | 0.007 – 0.249 |
|  | Leydig/Stroma cells | 0.097 | 0 – 0.200 |
|  | Peritubular cells | 0.330 | 0.145 – 0.568 |
|  | Sertoli cells | 0.490 | 0.334 – 0.776 |

### Isotretinoin stimulus

The expression of stimulated by retinoic acid-8 (*Stra8*) and promyelocytic leukemia zinc finger (*Plzf,* also referred to as *Zbtb16*) in neonatal testis CIVMs derived from PND 5 tissue (Figure 3.A for *Stra8* and 3.B for *Plzf*) and PND 10 tissue (Figure 3.C for *Stra8* and 3.D for *Plzf*) was quantified using RT-qPCR after exposure to isotretinoin for 24 hours. In the PND 5 derived cultures, Stra8 was not detectable in all samples from the CIVMs in the control, 0.3, 3, and 30nM dose groups. Samples with *Stra8* expression below the threshold of quantification, considered any sample with a cycle threshold of 37 or greater in the RT-qPCR analyses, are displayed as “X” in the figure. All CIVMs measured had quantifiable expression of Stra8 after exposure to the 300nM isotretinoin dose. The expression of *Stra8* was significantly higher in 300nM dose group relative to the control using a linear mixed effects model of delta-CT as a function of dose with a random intercept set for each biological sample to estimate marginal means, which were adjusted post-hoc using Tukey’s correction. Expression of *Plzf* was quantifiable in all PND 5 derived CIVMs across isotretinoin doses and the expression of *Plzf* was significantly lower in the 300nM dose group using the same analyses as reported for *Stra8*.

**Figure 3.**
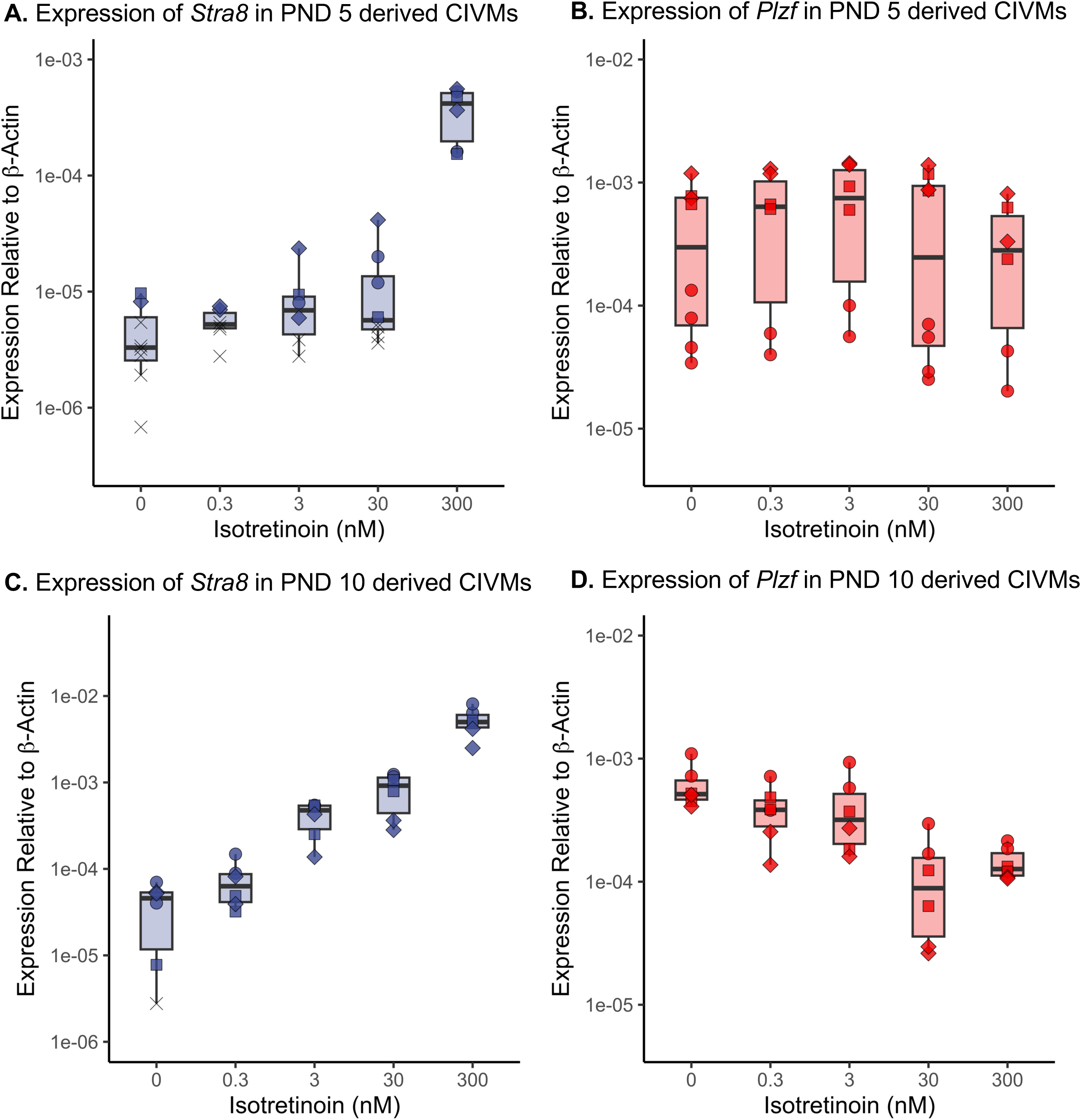
Expression of markers of spermatogonial phenotype after exposure to isotretinoin *in vitro.* CIVMs derived from neonatal testis from PND 5 and PND 10 testis were exposure to isotretinoin for 24 hours. (A) Expression of *Stra8* in PND 5 derived CIVMs determined by RT-qPCR. In the control, 0.3, 3, and 30 nM isotretinoin dose groups the expression of *Stra8* contained samples below the threshold of quantification. The expression of *Stra8* was significantly increased in the 300 nM isotretinoin dose group (p-value <0.0001). (B) Expression of *Plzf* in PND 5 derived CIVMs determined by RT-qPCR. All samples were above the threshold of quantification. The expression of *Plzf* in the 300 nM isotretinoin dose group was significantly lower than control (p-value of 0.026). (C) Expression of *Stra8* in PND 10 derived CIVMs determined by RT-qPCR. Aside from one control sample, all samples were above the threshold of quantification. The expression of *Stra8* was significantly increased in the 3,30, and 300 nM isotretinoin dose groups (p-values all <0.0001). (D) Expression of *Plzf* in PND 10 derived CIVMs determined by RT-qPCR. All samples were above the threshold of quantification. The expression of *Plzf* in the 30 and 300nM isotretinoin dose groups were significantly lower than control (p-values <0.0001).

In the PND 10 testis derived CIVMs, the expression of *Stra8* was detectable in all but one sample in control group. The addition of isotretinoin significantly increased the expression of *Stra8* in the 3, 30, and 300 nM PND 10 CIVM dose groups (p-values <0.001), with a trend in the 0.3 nM dose group (p-value of 0.10). The expression of *Plzf* was quantifiable in all PND 10 CIVMs, with a significant decrease in the 30 and 300 nM isotretinoin dose groups. Statistical significance in the PND 10 CIVMs was determined following the same approach as for the PND 5 CIVMs. In all cases, when the cycle threshold from RT-qPCR was below the threshold of quantification (CT of 37), the sample was marked as an X on the plot and CT value used for delta-CT calculations. If cycle threshold was below detection the sample CT was set to 40 for statistical calculations and plotting purposes. Data displayed as expression relative to beta-actin (housekeeping gene), by transforming the data to be 2 raised to the difference of the gene of interest and housekeeping gene (2^ [gene – HK]). This representation preserves the non-quantifiable samples, which are important for contextualizing the *Stra8* induction, and does not inflate fold-changes due to very lowly expressed genes. Due to the low number of non-detects in the PND 10 CIVM data, the data is also presented as log2 fold change (Supplemental Figure 1).

Expression of STRA8 in the PND 10 CIVM was confirmed through immunocytochemistry (supplemental figure 1). STRA8 protein expression was nuclear located and qualitatively at higher frequency in the PND 10 CIVM after 48 hours of 30 nM isotretinoin treatment compared to control.

### Retinol metabolism

The expression of *Stra8* in PND 10 testis CIVMs after addition of 3, 30, and 300 nM of retinol was determined through RT-qPCR (Figure 4.A). The addition of retinol to the CIVM increased *Stra8* expression across all doses (p-values <0.0001). The co-treatment of the CIVMs with retinol and 1 µM BMS-189453 resulted in no meaningful change in *Stra8* expression at 3 nM retinol, but significant decreases in *Stra8* expression at the 30 and 300 nM doses (p-values <0.0001). A two-hour pretreatment and continued co-exposure of retinol with 1 µM WIN-18446 resulted in a significant decrease in *Stra8* expression across all retinol doses (at 3 nM retinol a p-value <0.0001, at 30nM retinol a p-value of 0.0009, and at 300 nM retinol a p-value of 0.0142). The magnitude of *Stra8* reduction was relatively consistent in WIN-18446 pretreated CIVMs, where the differences in delta CT estimates were 1.67, 1.19, and 0.90 for 3, 30, and 300nM retinol, respectively. Changes in *Stra8* expression with BMS-189453 co-exposure were not consistent across retinol doses, where there was no significant difference in delta CT estimates at the 3 nM retinol dose, but estimated delta CT differences of 1.58 and 2.47 at 30 nM and 300 nM retinol respectively (calculated from estimated marginal means). Data displayed as log2 fold change relative to median of controls. The physiological relevance of retinoid metabolism within the PND 10 neonatal testis CIVM is additionally supported by detection of retinoid metabolism related genes within the CIVMs after three days *in vitro* (Figure 4.B).

**Figure 4.**
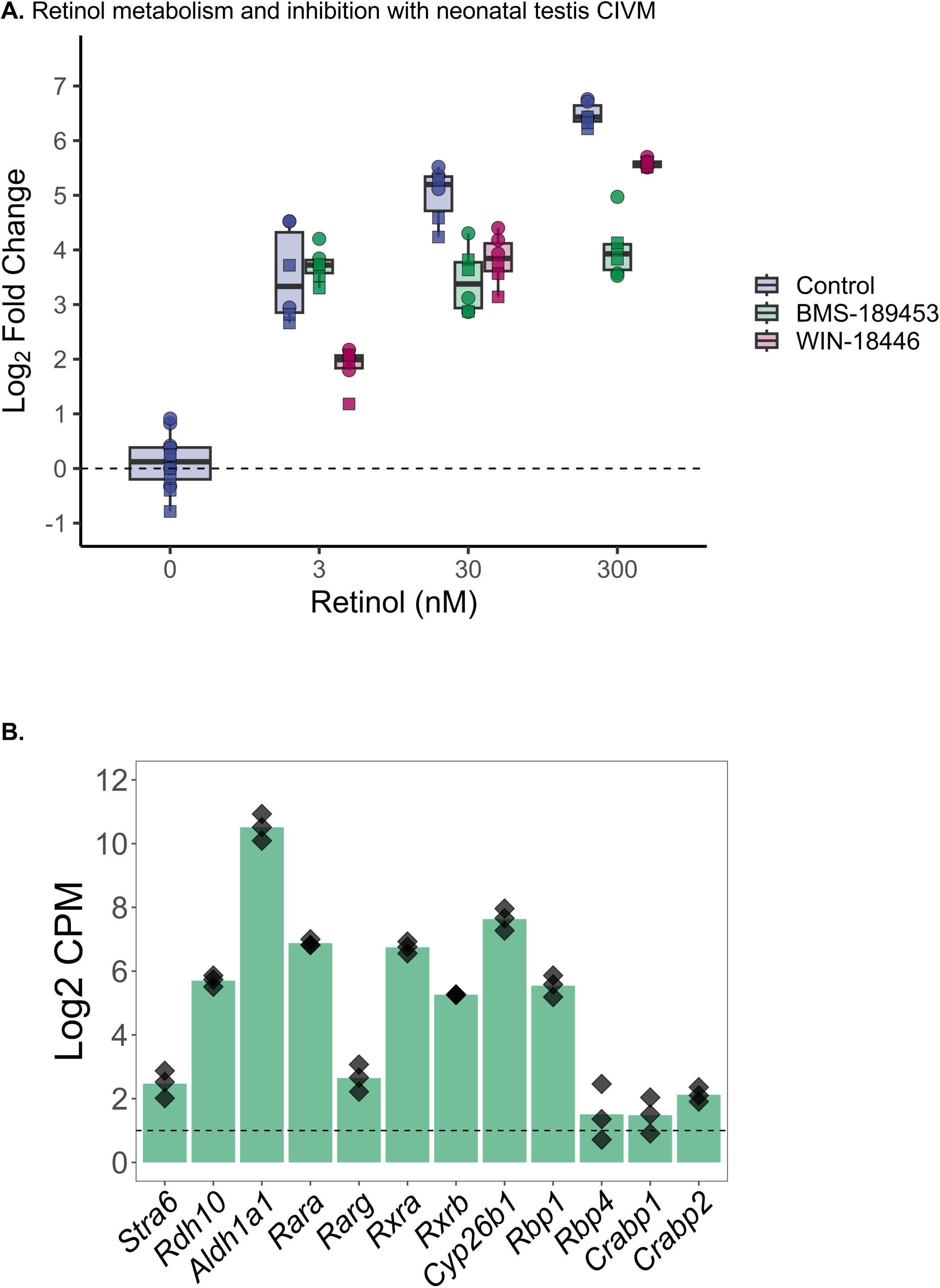
Retinol metabolism and inhibition within a PND 10 testis CIVM. Quantification of Stra8 activation after addition of retinol to PND 10 testis CIVM using RT-qPCR. (A) Addition of retinol into PND 10 testis CIVM at 3, 30, and 300 nM increases *Stra8* expression (blue). Co-exposure with BMS-189453 (green) and pre-treatment with WIN-18446 (red) show reduction in *Stra8* expression across doses. (B) Expression of key retinoid metabolism genes in RNA-seq analysis of PND 10 derived testis CIVMs. Includes expression of retinol transporter *Stra6*; metabolic enzymes *Rdh10*, *Aldh1a1*, and *Cyp26b1*; retinoid nuclear receptors and co-receptors *Rxr* (alpha and beta isoforms) and *Rar* (alpha and gamma isoforms); and transport proteins *Rbp4*, *Crabp1*, and *Crabp2*. Dashed line marks a log2 CPM of 1 as a gene expression threshold.

### Media formulation impact on spermatogonial phenotype

The impact of media supplementations on spermatogonial phenotype was measured through RT-qPCR to determine the expression of two marker genes, *Plzf* and *Star8,* over 15 days *in vitro*. Similar to the isotretinoin stimulation experiments (Figure 3), the variation in response was higher in the PND 5 derived testis CIVMs than in the PND 10 testis CIVMs. (A) The expression of *Plzf* in PND 5 and PND 10 CIVMs was quantified after 4 and 15 days *in vitro* maintained in base media (serum-free) conditions and after supplementation with SSC growth factors (GDNF, GFRA1, FGF2), lipid rich albumin (AlbuMAX), and defined bovine serum (Knockout Serum Replacement). In PND 5 testis CIVMs, the expression of *Plzf* was not statistically different with supplementation of SSC growth factors but was significantly decreased after four days of supplementation with lipid rich albumin (p-value of 0.0001) and defined bovine serum (p-value <0.0001) (Figure 5.A). Similarly, the expression of *Stra8* was not statistically different with supplementation of SSC growth factors but was significantly increased after four days of supplementation with lipid rich albumin and defined bovine serum (p-values <0.0001) (Figure 5.B). The expression of *Stra8* in PND 5 control CIVMs after four days *in vitro* was close to the limit of quantification, with many samples below quantification (a cycle threshold of 37). After 15 days *in vitro*, a qualitatively higher proportion of samples were above quantification, though no statistical assessment is reported due to high proportion of sample below quantification. The expression of Stra8 increased between days *in vitro* 4 and 15 with lipid-rich albumin supplementation (p-value 0.004) and decreased with defined serum supplementation (p-value <0.0001). The expression of *Plzf* significantly decreased between days *in vitro* 4 and 15 in the control media CIVMs (p-value <0.0001), while all supplemented CIVMs had no notable change in *Plzf* expression (Figure 5.A).

**Figure 5.**
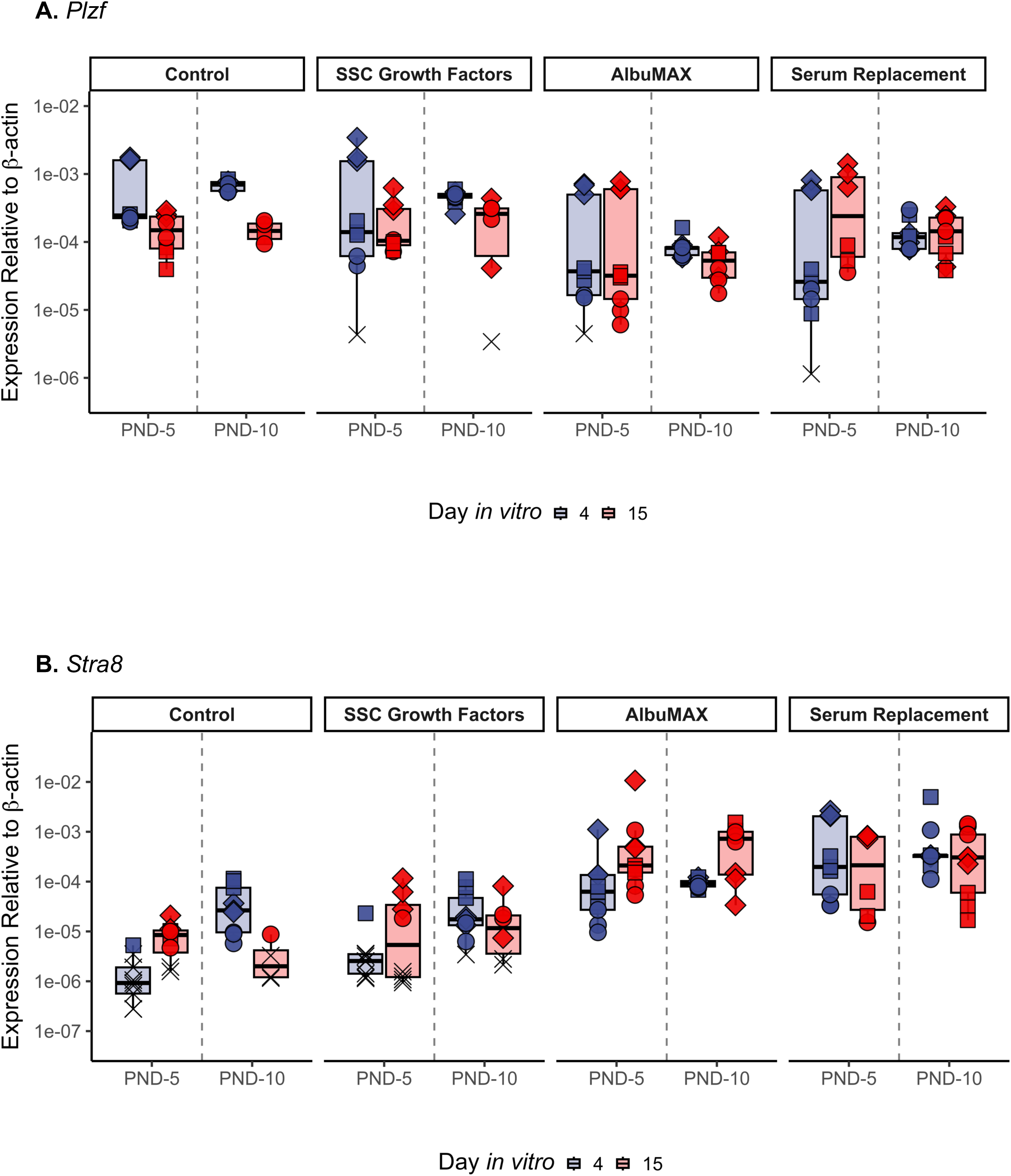
Spermatogonial phenotype after media supplementation media for 15 days *in vitro.* The expression of *Plzf* and *Stra8* quantified by RT-qPCR at day *in vitro* 4 and day *in vitro* 15 in PND 5 tissue CIVMs and PND 10 tissue CIVMs in control and media supplementation conditions. Color indicates day *in vitro*, dashed line divides plot by PND 5 CIVMs and PND 10 CIVMs. Points that are “X” indicate samples that were below the limit of quantification (cycle threshold of 37). Data displayed as expression relative to beta-actin (housekeeping gene (HK), (2^ [gene – HK])).

In PND 10 testis tissue derived CIVMs, the expression of *Plzf* trended lower, but was not statistically different with supplementation of SSC growth factors (p-value of 0.07). *Plzf* was significantly decreased after four days of supplementation with lipid rich albumin and defined bovine serum (p-values <0.0001) (Figure 5.A). *Stra8* expression was not statistically different with supplementation of SSC growth factors but was significantly increased after four days of supplementation with lipid rich albumin (p value of 0.011) and defined bovine serum (p-values <0.0001) (Figure 5.B). After 15 days *in vitro*, a higher proportion of CIVMs maintained in control media or SSC growth factors had *Stra8* expression below quantification relative to day *in vitro* four. The expression of *Stra8* increased between days *in vitro* 4 and 15 with lipid-rich albumin supplementation (p-value 0.03) but was unchanged with defined serum supplementation (p-value 0.96).

Overall, the culture viability was not qualitatively different with media supplementation and was qualitatively similar to our previous investigations using similar culture methods [19]. Notably, media supplementations that contained serum products (Knockout Serum Replacement and AlbuMAX; Figures 6.A-D) led to distinct morphological differences relative to the serum-free cultures after four days (Control and SSC growth factors supplementation; Figures 6.E-H) in CIVMs derived from both developmental stages (PND 10 [Figure 6.C-D,G-H] and PND 5 [Figure 6.A-B,E-F]) after four days *in vitro*. Sertoli cells and spermatogonia were retained through 15 days *in vitro* and confirmed through immunocytochemistry for SOX9 (Sertoli cells) and DDX4 (spermatogonia) (Supplemental Figures 2.A-B).

**Figure 6.**
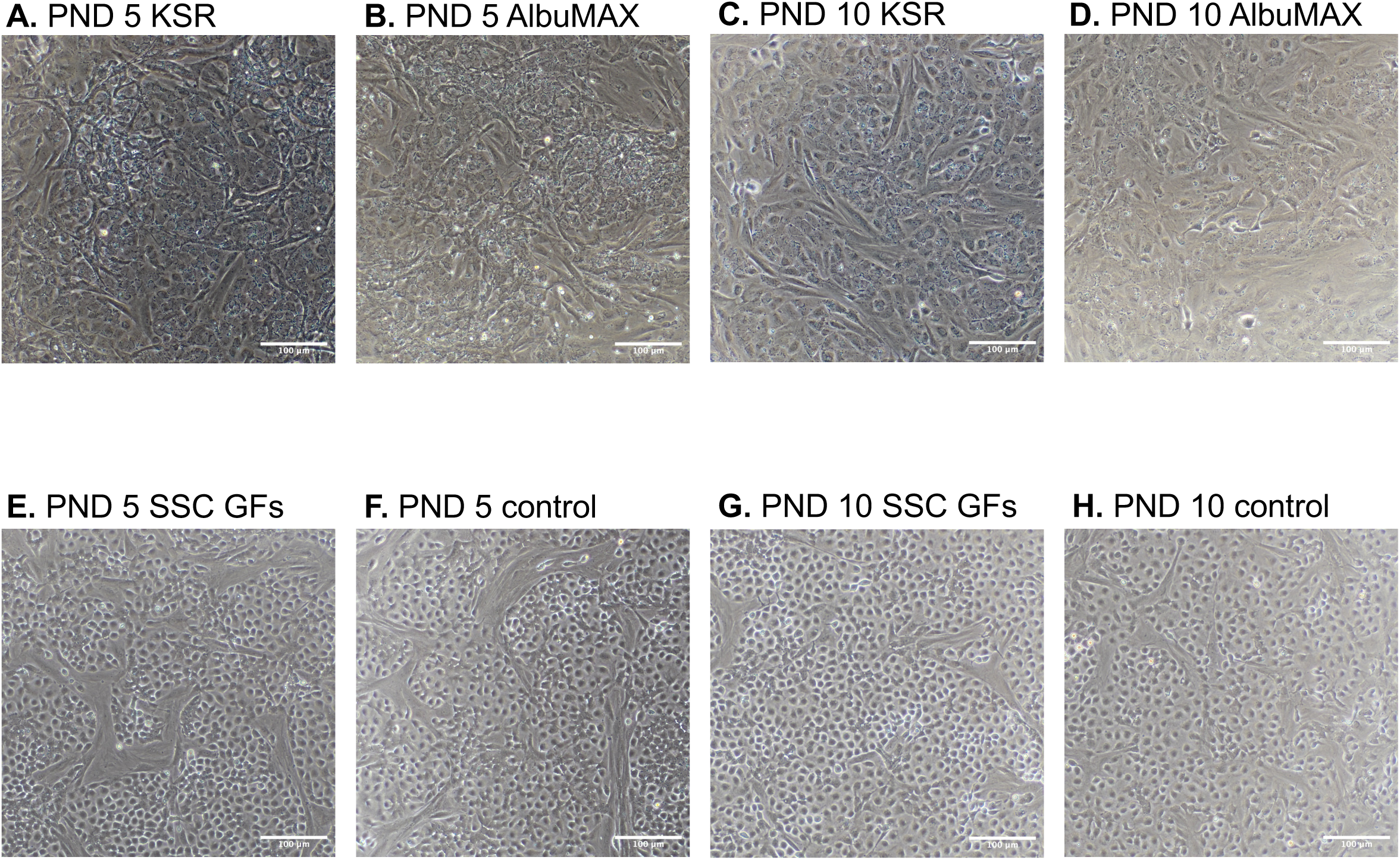
Media supplementation cell morphology. Brightfield images of PND 5 testis tissue derived CIVMs (A-B, E-F) and PND 10 testis tissue derived CIVMs (C-D, G-H) after 4 days *in vitro* maintained in either: control media, SSC growth factors (GDNF, GFRA1, FGF2), lipid rich albumin (AlbuMAX), and defined bovine serum (Knockout Serum Replacement).

## Discussion

Neonatal rodents remain the preeminent model organism for studying testicular development and reproductive dynamics. The extension of *in vivo* rodent research to *in vitro* models provides a reductionist approach to expand the breadth of molecular analyses and biological manipulation possible in reproductive and developmental studies. Further, the progression of *in vitro* models based on non-human tissue remains a critical way to reduce the need for animals in toxicology and preclinical research for organ systems where human tissue is not available for research, like the developing testis [37]. However, *in vitro* models require careful assessment to identify biological significance from *in vitro* artifacts and to benchmark model phenotype within a dynamic developing system. The impact of *in vitro* culture on spermatogonial phenotype is a critical consideration for a range of reproductive and developmental research: from novel models of *in vitro* spermatogenesis to molecular analyses of toxicants on specific spermatogonial stages. We find that while complex *in vitro* models (CIVMs) seeded with primary testis cells from post-natal day (PND) 5 and PND 10 Sprague Dawley rats retain similar cell type compositions, with transcriptomic signatures of Sertoli cells, Leydig cells, Germ cells, and peritubular myoid cells, there are distinct differences in the overall transcriptome, transcription factor activity, and retinoid responsiveness. These manifest in disparate responses of spermatogonia within the CIVMs at PND 10 to those at PND 5 after exposure to physiologically relevant levels of retinoid and distinct responses to common cell culture media supplements. These findings have critical implications for in vitro studies of spermatogonial dynamics, where various neonatal ages or media supplements have been used. Our results support that in neonatal rat models, PND 10 testis can be a consistent and reproducible model for retinoic acid metabolism and spermatogonial response.

### Transcriptomics analyses

Overall, the transcriptome of CIVMs from PND 5 rat testis was distinct from that of CIVMs seeded from PND 10 rat testis. We found that more than 700 genes were differentially expressed between CIVMs at the two developmental stages following the standard edgeR workflow. To apply biological meaning to these differentially expressed genes, we considered if certain annotated pathways were over-represented within our differentially expressed gene set, suggesting those pathways may be specifically different between the developmental stages. This over-representation analysis revealed significant enrichment in a number of pathways (supplemental table 1), but review of this list identified specific developmentally relevant pathways pertaining to retinoic acid response and metabolism (Table 1) that were differentially expressed between the PND 5 and PND 10 CIVMs. Assessment of the major genes considered part of retinoic acid metabolism in the testis revealed that while certain retinoic acid metabolism related genes were differentially expressed (i.e. *Stra6, Cyp26b1, Rbp1, Crabp2; adj. p-value <0.05*), each was upregulated in PND 5 CIVMs relative to the PND 10 CIVMs (supplemental figure 6). Aside from *Cyp26b1*, which removes the bioactive retinoic acid and is critical for normal testis development [38], the impact of the differential expression of *Stra6*, *Rbp1*, or *Crabp2* is uncertain as these are responsible for intracellular transport or binding of retinoids. This follows the trend of the broader dataset where the majority of genes differentially expressed within the retinoic acid related ontologies (GO:0042573, GO:0071300, and GO:0001523) are upregulated in PND 5 relative to PND 10 (supplemental figure 7). To further address the differences in transcriptome between the PND 5 and PND 10 testis CIVMs, the differentially expressed genes were modeled through their canonical association with transcription factors (Figure 1.D). Target genes of the transcription factor cleraxis (*Scx),* associated with Sertoli cell growth and maturation [39], were more active in the PND 5 CIVMs than in the PND 10 CIVMs. Limited information is available on the role of *Scx* in the developing testis. Interestingly, the previous report on *Scx* in the testis [39], using freshly isolated rat Sertoli cells, identified a transient increase in *Scx* expression at PND 10, whereas our report did not find a significance difference in expression of *Scx* itself between PND 5 and PND 10 CIVMs, rather an increase in inferred *Scx* activity in PND 5 CIVMs based on differential expression of target genes. The canonical Sertoli cell marker, *Sox9*, was not differentially expressed between the CIVM developmental stages, though the transcription factor activity inference identified that multiple Sry-type HMG box proteins (SOX Family), *Sox9*, *Sox5*, and *Sox6,* were more active in the PND 5 CIVMs. Based on previous reports (referenced in [40]), it is unlikely that all of these SOX transcription factors were expressed in the same cell type, however, they are related to a coordinated response with *Col2a1* upregulation, which is significantly upregulated in the PND 5 CIVMs reported here. This supports a coordinated matrix remodeling in the PND 5 CIVMs after three days *in vitro* that is not present in the PND 10 CIVMs after three days *in vitro*.

### Deconvolution analyses

While there were significant differences in the transcriptomes of CIVMs derived from PND 5 and PND 10 testis, our deconvolution analyses support that there are not broad differences in cell type proportions between these developmental stages (Figure 2). Qualitatively, the whole-mount imaging of the primary seminiferous tubules confirms that DDX4+ germ cells are present at both developmental stages in the rat, though variation across tubule length make quantitative assessments unreliable (supplemental figure 2). This is also supported by a lack of significant diffntial expression of canonical somatic cell marker genes: *Cyp11a1* for Leydig cells, *Sox9* and *Wt1* for Sertoli cells, and *Acta2* for peritubular myoid cells (Figure 1.A). Our analyses find that the bulk transcriptome of both the PND 5 and PND 10 CIVMs are composed of ∼ 9-10% germ cells, ∼10-11% Leydig cells, ∼30-33% peritubular myoid cells, and ∼48-49% Sertoli cells (Table 2.). The PND 10 testis are larger than the PND 5 testis, though our data support that *in vitro* each maintains a similar cell type proportion over time *in vitro,* suggesting the cells are proliferating at similar rates. The convergence on this cell type proportion may be a function of differential viability in culture, where germ cells may be more fragile compared to peritubular cells which may be more robust, as well as the lack of pituitary signaling that can drive cell type specific proliferation. *In vivo*, it is expected that the Sertoli cells are highly proliferative at PND 5, whereas at PND 10 the spermatogonia and other somatic cells are highly proliferative as well [6].

### Isotretinoin stimulus

While the cell type proportions may be similar between the PND 5 and PND 10 CIVMs, the response to retinoic acid is distinctly different between the two CIVM developmental stages based on induction of stimulated by retinoic acid-8 (*Stra8,* a marker of spermatogonial differentiation) and promyelocytic leukemia zinc finger (*Plzf*; a marker of undifferentiated spermatogonia) expression (Figure 3). Using doses of isotretinoin that are reflective of physiologically relevant levels of bioactive retinoids based on published adult human tissue data [41], where 0.3 nM is reflective of isotretinoin in testicular homogenate, 3 nM is reflective of isotretinoin in serum, while 30nM and 300nM are dose escalations. Isotretinoin (13-cis retinoic acid), is an endogenous isoform of the bioactive all-trans-retinoic acid (ATRA) and is used as a pharmaceutical as it is a poor substrate for the CYP26B- enzymes that degrade bioactive retinoids [42]. In the PND 5 testis derived CIVMs, *Stra8* was largely undetectable at baseline or at the 0.3 nM dose of isotretinoin. In the 3 nM and 30 nM dose groups, there were more samples with quantifiable expression of *Stra8* (i.e., above the 37-cycle threshold in RT-qPCR), though only at the 300 nM dose group did *Stra8* expression increase significantly in the PND 5 CIVMs (Figure 3.A). In the same PND 5 CIVMs, *Plzf* was expressed above quantitation in all samples, but showed no significant dose-relationship (Figure 3.B). In the PND 10 CIVMs, there was a significant increase in *Stra8* across isotretinoin doses parallelling a decrease in *Plzf* expression (Figure 3.C-D). The response of spermatogonia in PND 10 CIVMs to physiological levels of bioactive retinoic acid (isotretinoin) is a promising advance in *in vitro* spermatogonial differentiation studies, where supraphysiological levels may exceed relevant enzyme capacity, decreasing translational power. Our finding that nanomolar doses of isotretinoin elicit a spermatogonial response suggest that a possible “target” for tissue concentrations of RA in patients with NOA in clinical trials of retinoid-based treatments of men being treated for infertility from reduced sperm production may be at least 30 nM or higher to initiate or stimulate spermatogenesis. While follow-up clinical studies to refine this hypothesis are needed, clinical dose ranges can be refined using this in vitro method. This is very advantageous for dosing high affinity interactions, like the retinoids and retinoid receptors, where the nanomolar range is sufficient for activity. Particularly in cases where a therapeutic may target increased spermatogonial differentiation as a treatment for a non-obstructive oligozoospermia, possible therapeutics could be trialed *in vitro*, where impacts on spermatogonial differentiation can be measured in a reductionist system at the cellular level.

While the response to exogenous isotretinoin is important for mechanistic studies of spermatogonial differentiation and for screening therapeutics to increase sperm count, the metabolism of bioactive ATRA itself is a critical step to recapitulate *in vitro* to progress *in vitro* investigation of infertility causes and screening potential contraceptives. Estimates show that a very small percentage (< 1%) of bioactive retinoic acid within the testis originated elsewhere and was brough via circulation, rather the extensive metabolic network of retinoid metabolizing enzymes are responsible for maintaining the intratesticular retinoid environment [43]. Inhibition of the aldehyde dehydrogenase enzymes (ALDH family) within the testis has historically been a focus for development of male contraceptives [44,45], and inhibiting retinoic acid activity continues to be a target for novel contraceptives [46–48]. Our results here show that the PND 10 testis derived CIVM retains the capacity to metabolize retinol to ATRA at levels sufficient to drive *Stra8* expression (Figure 4.A), our RNA-seq analyses show that genes for key the critical metabolizing enzymes, transporters, and binding proteins are expressed within the CIVMs (Figure 4.B). Through the addition of specific inhibitors of retinoic acid formation by inhibiting ALDH retinyl metabolism (WIN-18446) and retinoic acid receptor activation by allosterically binding the RAR complex (BMS-189453), our PND 10 CIVM data supports that these critical processes are retained *in vitro* and are inducible and inhibitable (Figure 4.A).

Given our data supporting that these developing testis CIVMs are responsive to retinoids, we investigated if residual retinoids in commonly used cell culture supplements were sufficient to drive spermatogonial phenotype. Using three media supplement approaches common for *in vitro* testis models: spermatogonial stem cell (SSC) growth factors [20], lipid-rich albumin (AlbuMAX, Thermo Fisher Scientific) [21], and a defined bovine serum (Knockout Serum Replacement) [22,23], we found that addition of serum derived components impacted the phenotype of the spermatogonia *in vitro* (Figure 5). Using *Plzf* and *Stra8* gene expression as metrics for undifferentiated and differentiating spermatogonia respectively, our data support that addition of serum derived components, lipid rich albumin and defined bovine serum, increase the expression of Stra8 after four days *in vitro* in both the PND 5 and PND 10 derived testis CIVMs. The upregulated of *Stra8* in the PND 10 CIVMs was accompanied by a significant decrease in *Plzf*, while there was not a significant decrease in *Plzf* expression in the PND 5 CIVMs (Figure 5). Notably, the additions of lipid rich albumin and defined bovine serum were sufficient to drive *Stra8* expression in the PND 5 CIVMs, while it was generally unquantifiable under serum-free (control and SSC growth factors) conditions. This is hypothesized to be related to residual retinoids within the serum products, though not directly measured in this report. Previous reports show that residual retinoids in cell culture products can be biologically active and our data presented in this report support that the activity of residual retinoids is critical to consider in *in vitro* testis models [49].

When these CIVMs were maintained for 15 days *in vitro*, there was a significant decrease in *Plzf* expression in the control media CIVMs derived from both ages, PND 5 and PND 10, there was not a significant decrease in *Plzf* gene expression in the SSC growth factor CIVMS between *in vitro* days 4 and 15. While 15 days was likely not a long enough culture period to see significant proliferation of spermatogonia, where previous studies are carried out for up 10 weeks and see significant expansion of SSCs [20], our data suggest the addition of SSC growth factors may support the undifferentiated spermatogonial phenotype (characterized here by *Plzf* expression). As expected based on the isotretinoin exposure experiments (Figure 3), there is little to no baseline expression of *Stra8* in PND 5 testis CIVMs, whereas it is consistently expressed above quantification in PND 10 derived CIVMs. Additionally, there is a lower variation of spermatogonial response within the PND 10 derived testis compared to a much larger variation in the PND 5 derived CIVMs. This is an interesting finding considering a previous work which found that retinoic acid responsiveness was a factor of spermatogonial phenotype, as KIT+ differentiating spermatogonia were less responsive. However, this could be due to comparing relative changes, where the expression of *Stra8* in already differentiating spermatogonia is less inducible than undifferentiated spermatogonia [50].

Overall, we propose that *in vitro* models of the developing testis are powerful tools to study the dynamics of retinoids and spermatogonial phenotype. We find that physiological levels of retinoids are sufficient to drive *Stra8* expression in testis CIVMs derived from PND 10 tissue. We report similarities in cell type proportions, but distinct differences in transcriptomes and inferred transcription factor activity of CIVMs derived from PND 5 testis and PND 10 testis. Additional research is warranted to further investigate the dynamics of spermatogonia *in vitro*. A model that recapitulates the spermatogonial niche dynamics that support proliferation of undifferentiated spermatogonia through retinoic acid driven differentiation will advance reproductive biology and toxicology and reduce the need for lengthy *in vivo* studies.

## Data Availability

All code is available on GitHub: github.com/bchansen3/Hansen_2025_Spermatogonia

RNA-seq data is available as FASTQ files and gene counts on the Gene Expression Omnibus: GSE314458.

Any other additional data available upon request.

## Author Contributions

Conceptualization: BCH, EMF, JKA, EJK; Methodology: BCH, SMH, LAH; Formal analysis: BCH; Resources: Z.J; Writing—original draft: BCH; Writing—review & editing: BH, SMH, LAH, EMF, JKA, EJK; Supervision: JKA, EMF, EJK; Funding acquisition: BCH, EMF, EJK.

## Acknowledgements

The authors would like to thank members of the Isoherranen laboratory at the University of Washington for their expertise on using retinoids *in vitro*, specifically: Aprajita Yadav, Aurora Authement, and Jiayao Chen.

The authors would like to thank Dashiel Cockrill for insightful questions and support on the experiments presented here.

The authors would also like to thank Dr. Dale Hailey at the UW ISCRM Garvey Imaging core for microscopy support and the University of Washington Department of Environmental and Occupational Health Sciences IT team for maintenance and support of computing servers.

## Supplemental Tables

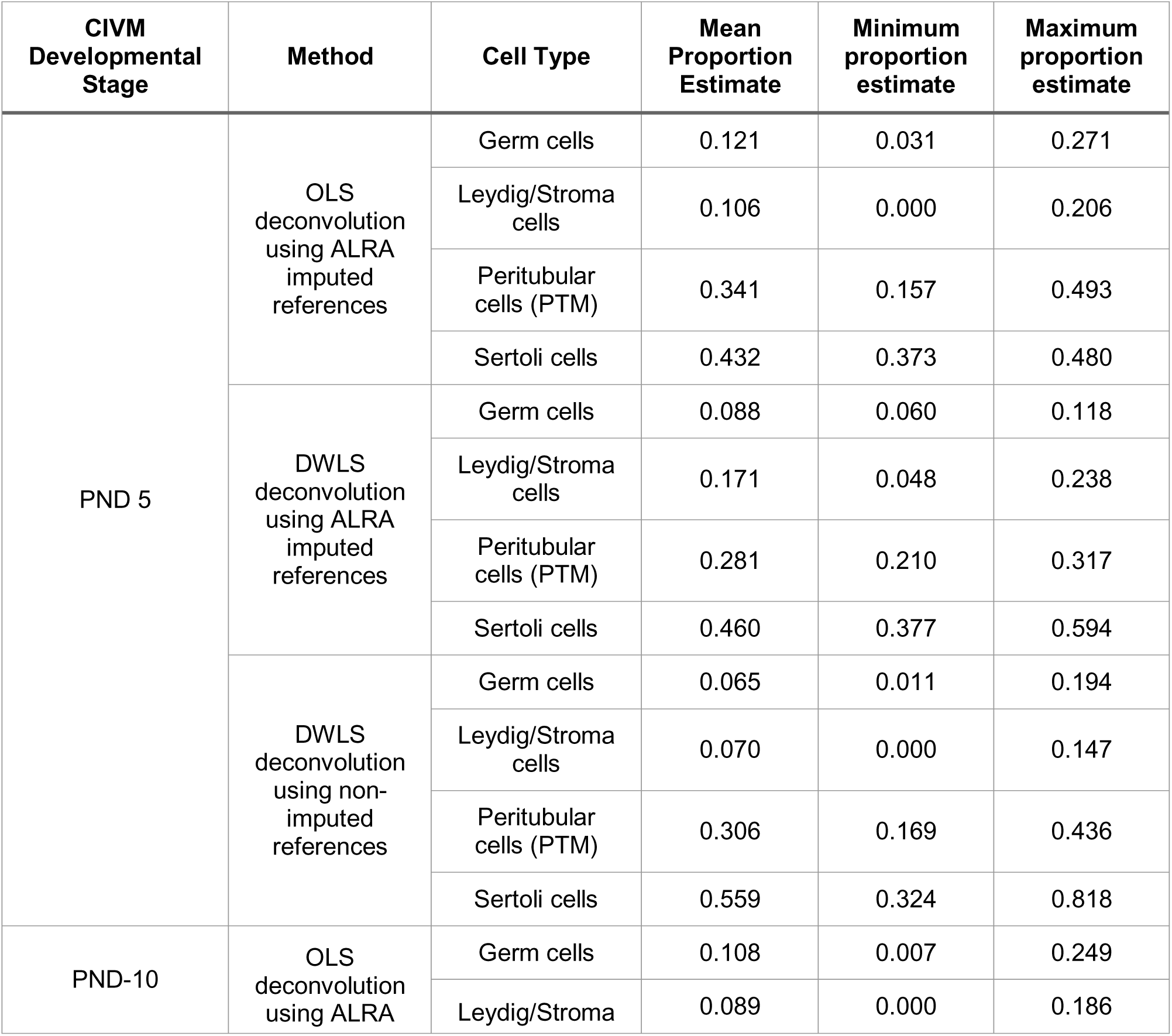
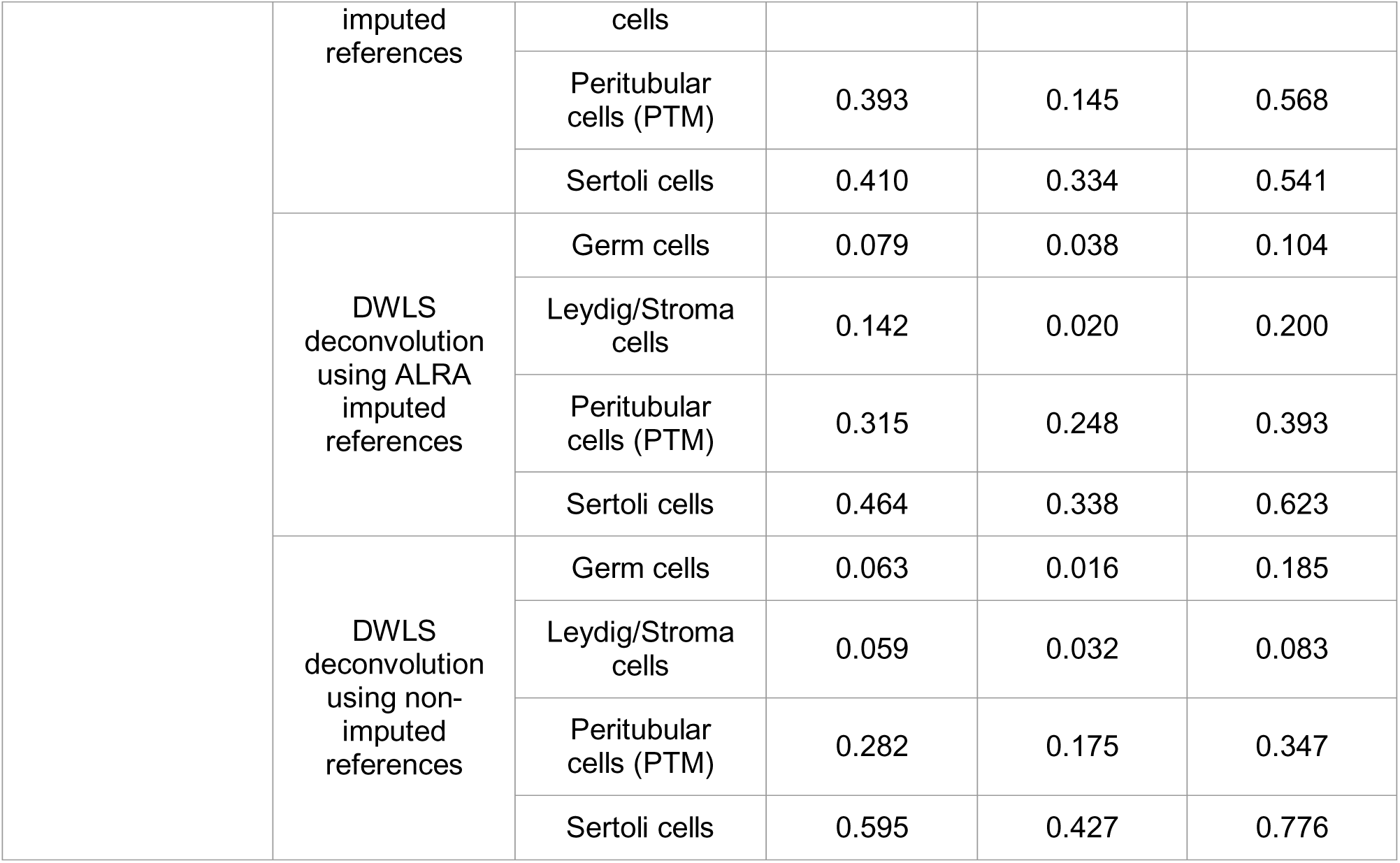
Supplemental Table 1

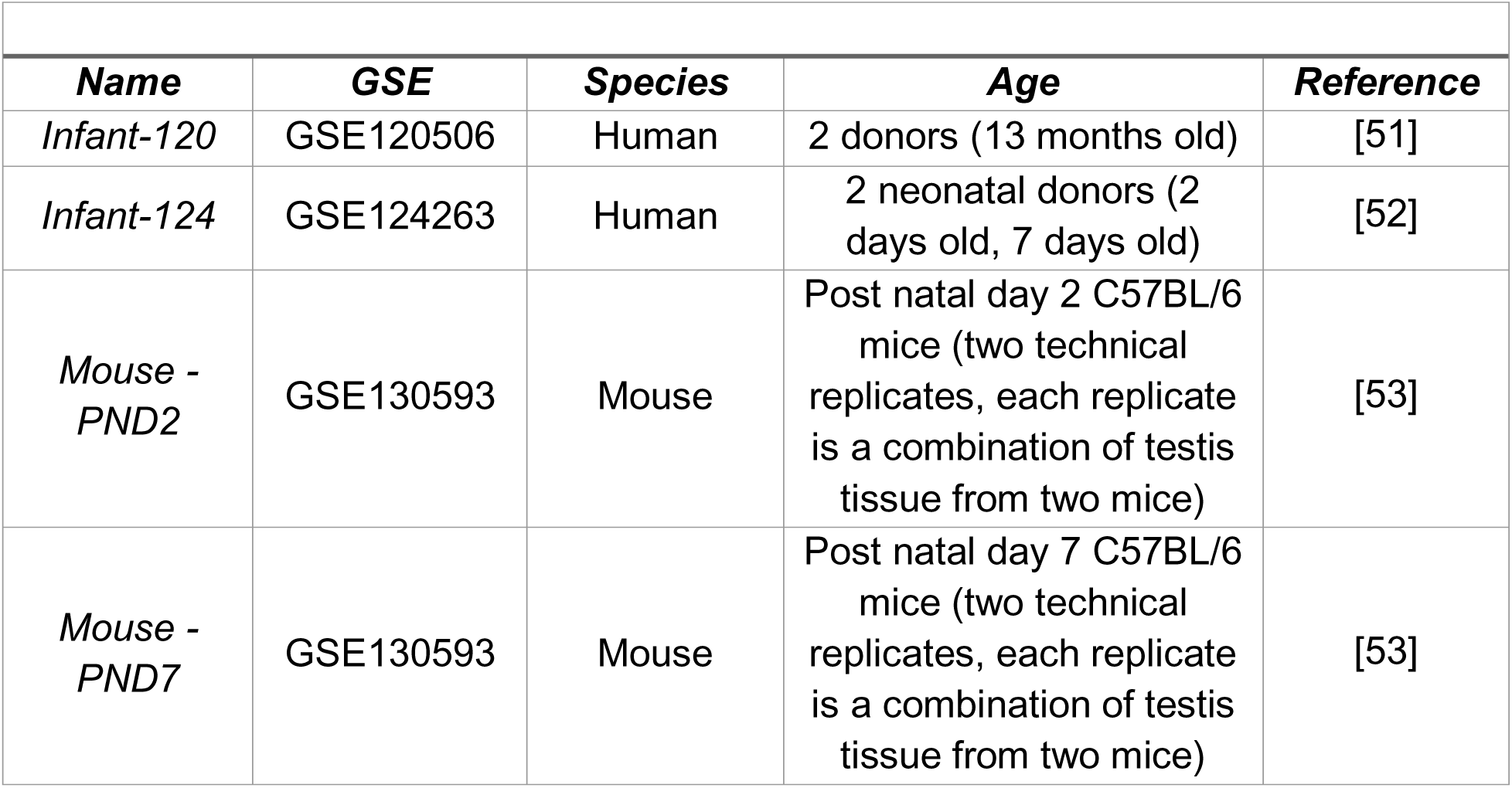
Supplemental Table 2

Supplemental Table 2
| **Name** | **GSE** | **Species** | **Age** | **Reference** |
| --- | --- | --- | --- | --- |
| *Infant-120* | GSE120506 | Human | 2 donors (13 months old) | [51] |
| *Infant-124* | GSE124263 | Human | 2 neonatal donors (2 days old, 7 days old) | [52] |
| *Mouse - PND2* | GSE130593 | Mouse | Post natal day 2 C57BL/6 mice (two technical replicates, each replicate is a combination of testis tissue from two mice) | [53] |
| *Mouse - PND7* | GSE130593 | Mouse | Post natal day 7 C57BL/6 mice (two technical replicates, each replicate is a combination of testis tissue from two mice) | [53] |

