## Supplemental Figure 1 for "Differential retinoic acid responses across testicular development *in vitro*"

**Supplemental Figure 1. Immunocytochemistry of STRA8 expression in PND 10 CIVMs after 48 hours of exposure to 30nM isotretinoin.**

**A.**

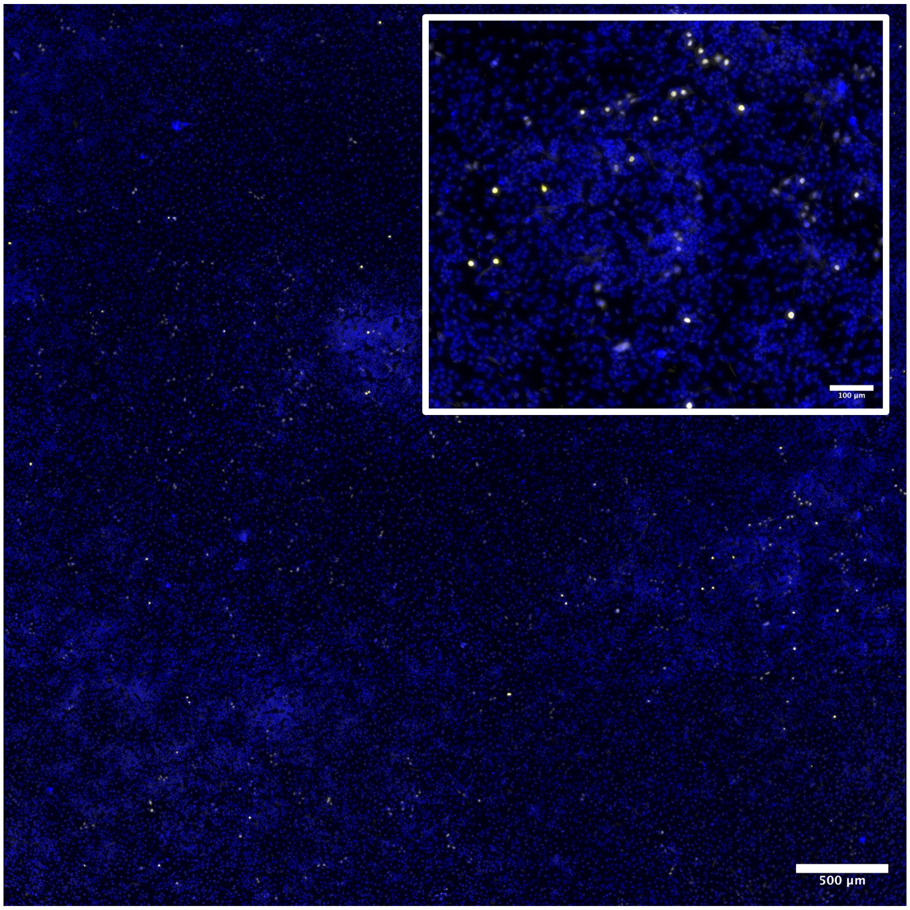


**B.**

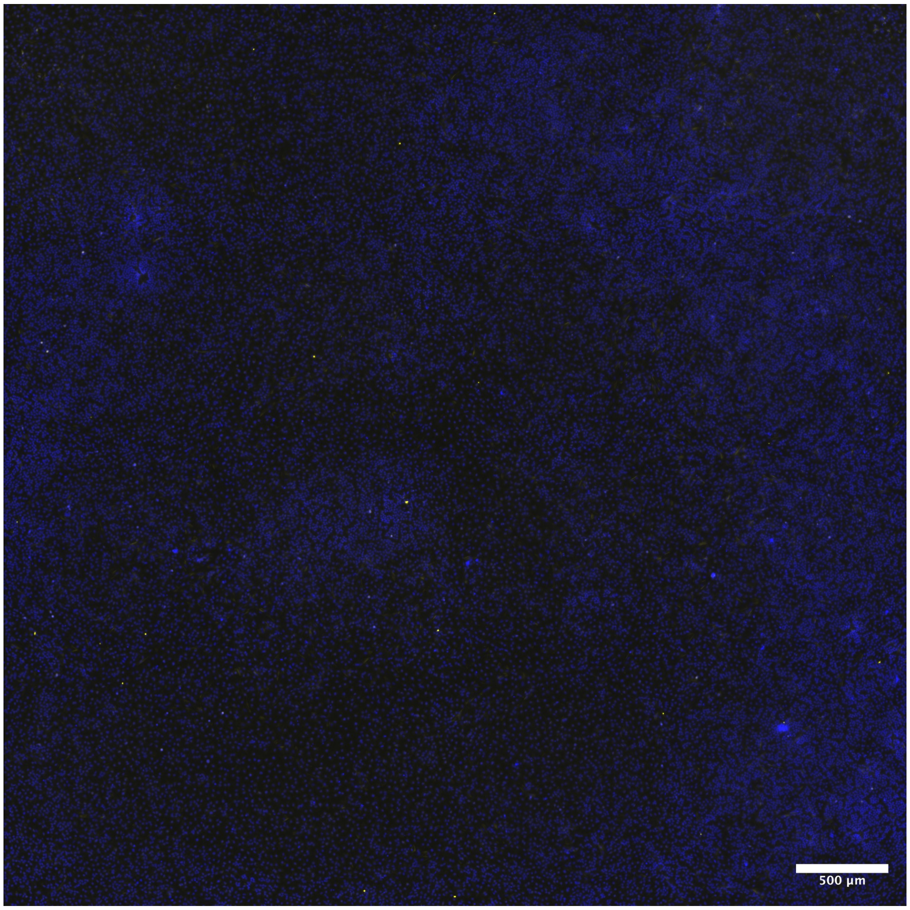
