## Supplementary figures and images for "Differential retinoic acid responses across testicular development *in vitro*"

### Supplemental Figure 2

**Supplemental Figure 2.** Immunocytochemistry

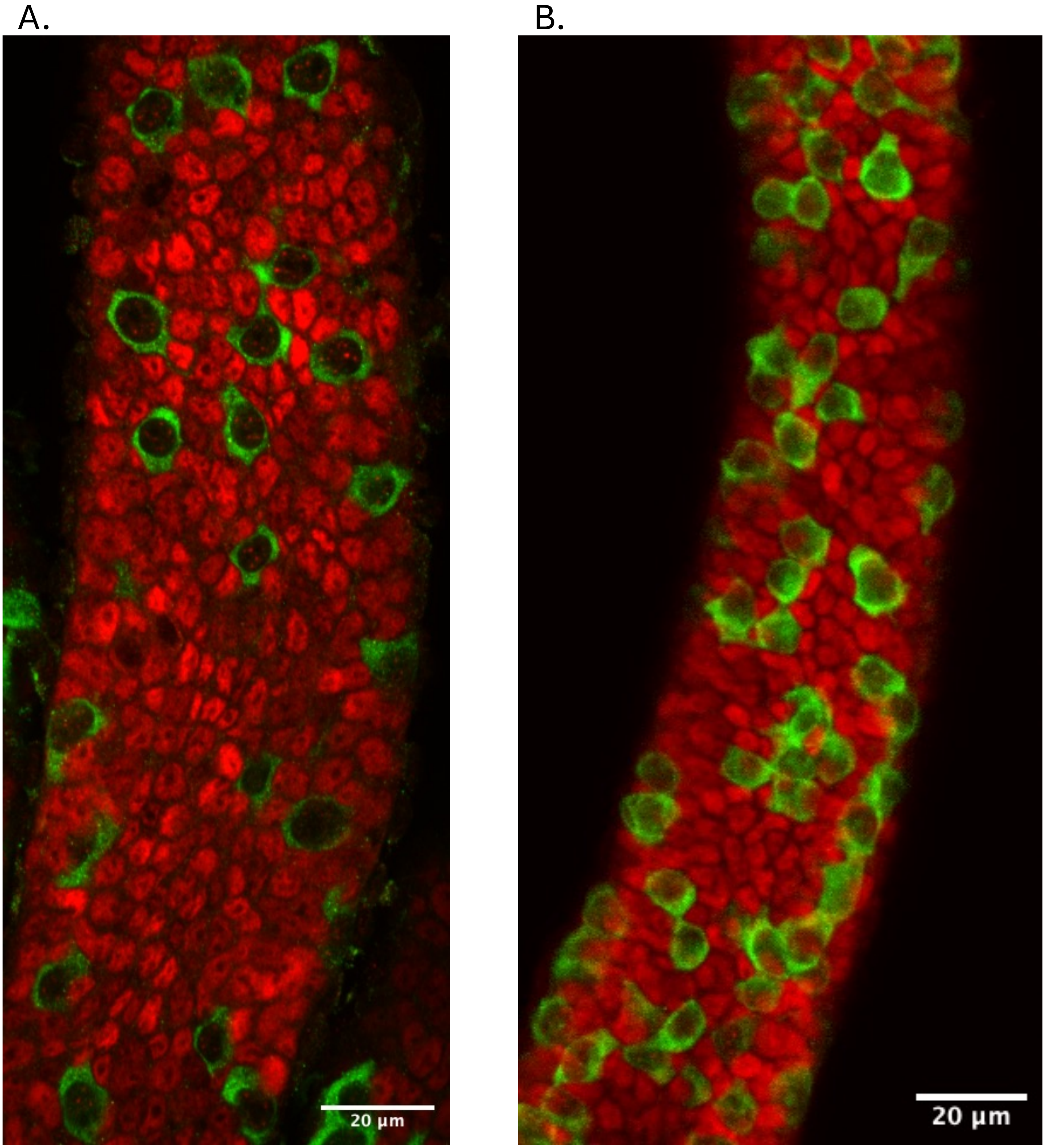

### Supplemental Figure 3

**S3. Widefield fluorescent immunocytochemistry of PND 5 and PND 10 CIVMs.**

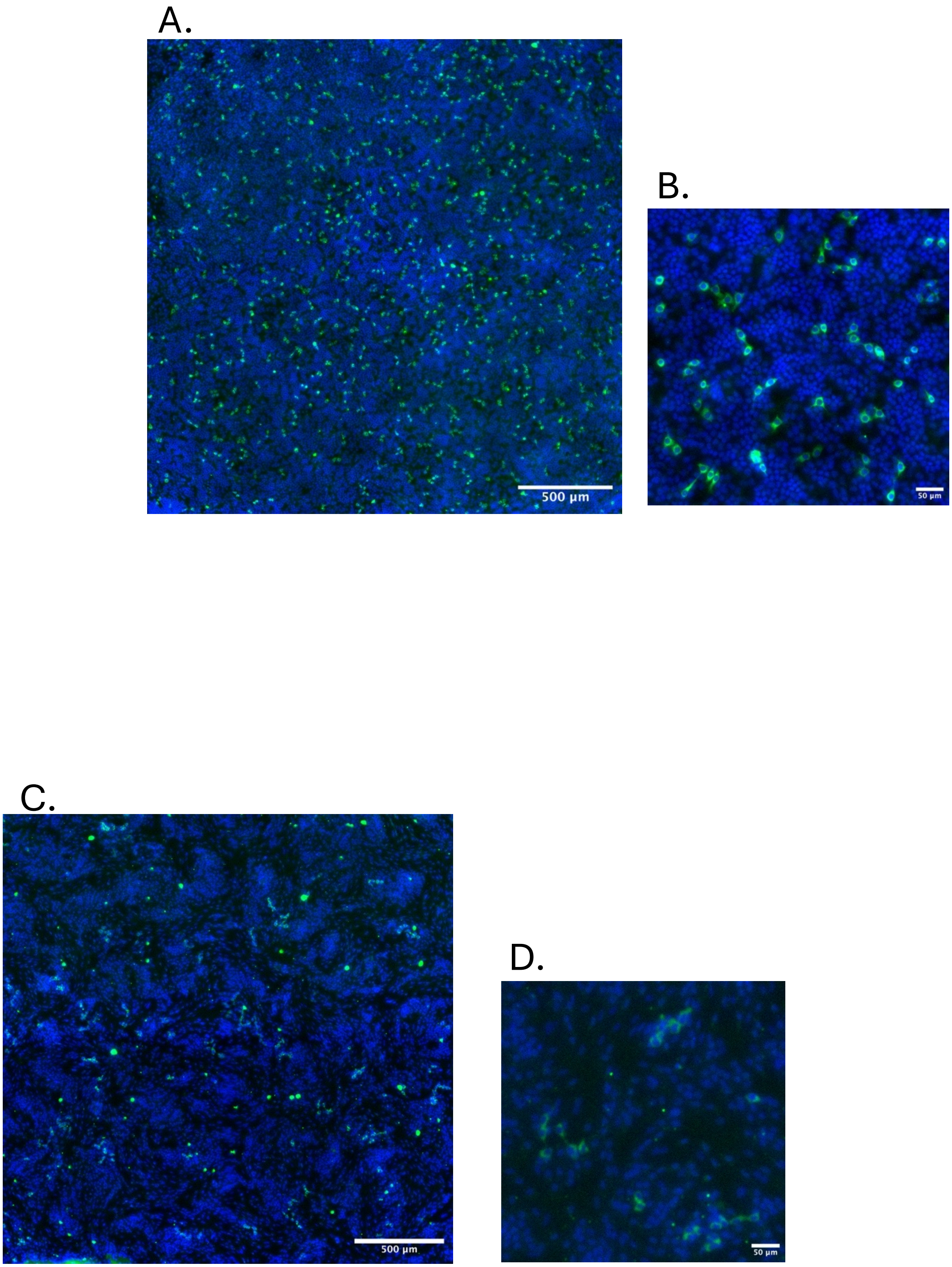

### Supplemental Figure 7

**Supplemental Figure 7. Retinol metabolism and inhibition within PND 10 and PND 5 testis CIVMs.**

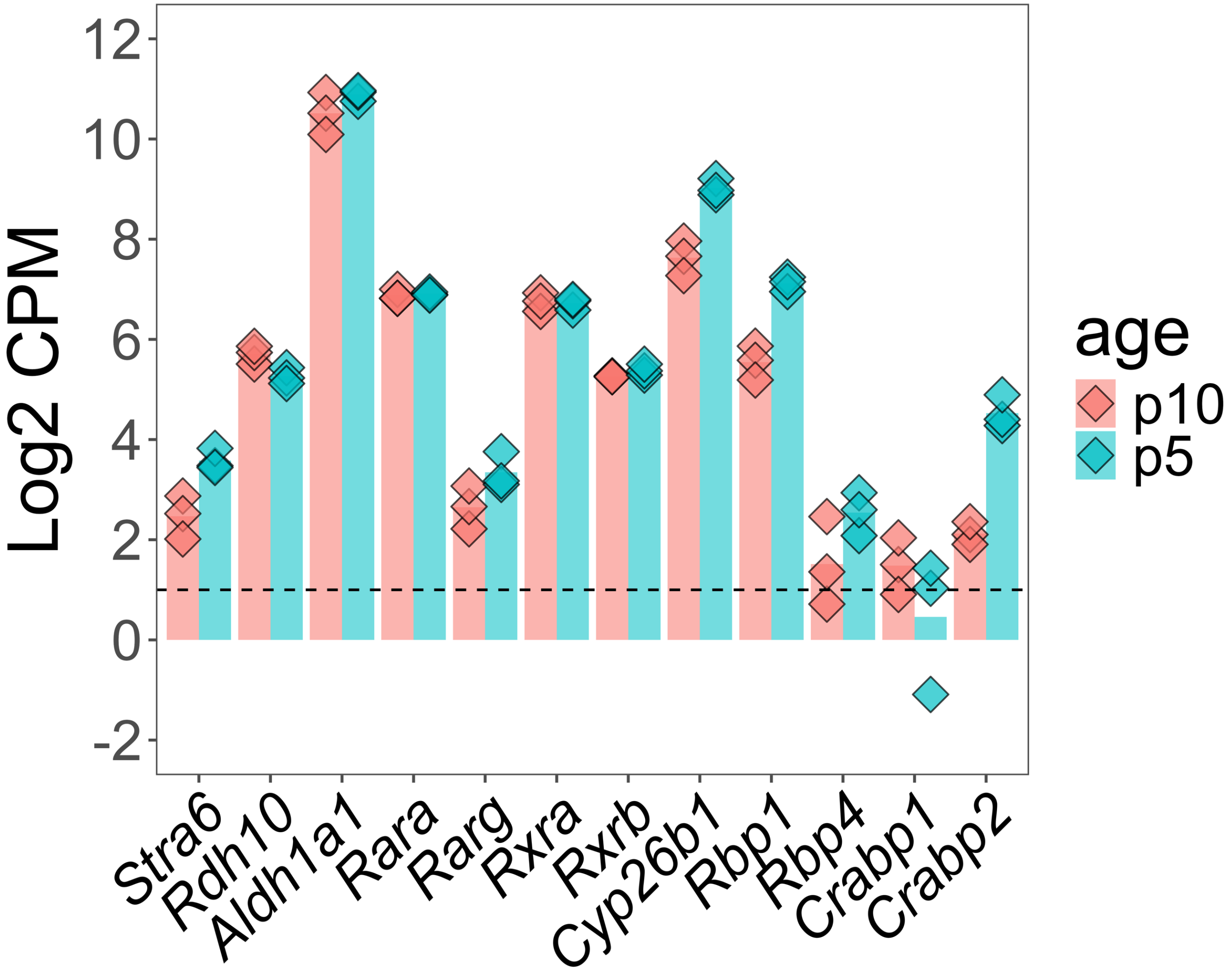
