## Supplemental Figure 4 for "Differential retinoic acid responses across testicular development *in vitro*"

**Supplemental Figure 4. Log2 fold change of markers of spermatogonial phenotype after exposure to isotretinoin in vitro.**

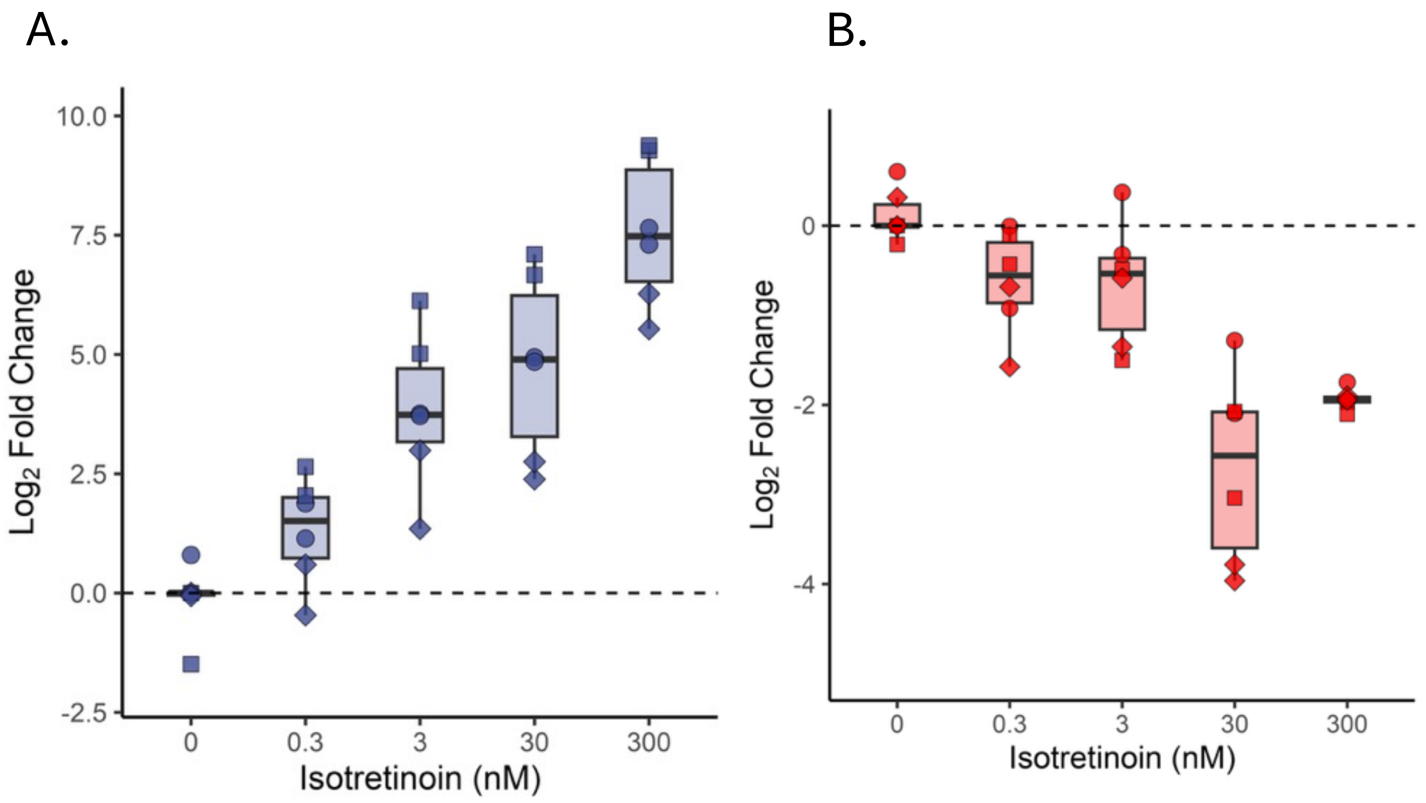
