## Supplemental Figure 5 for "Differential retinoic acid responses across testicular development *in vitro*"

**Supplemental Figure 5. Fluorescent immunocytochemistry of example PND 5 and PND 10 CIVMs after 15 days in vitro.**

A.

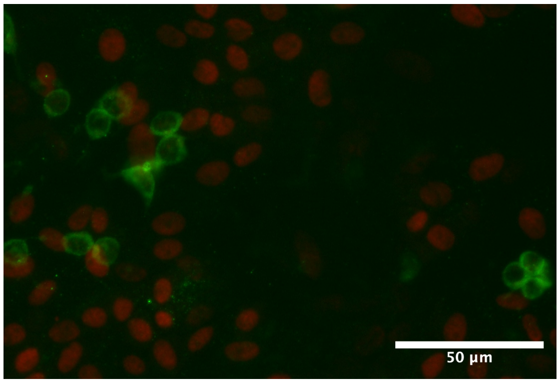


B.

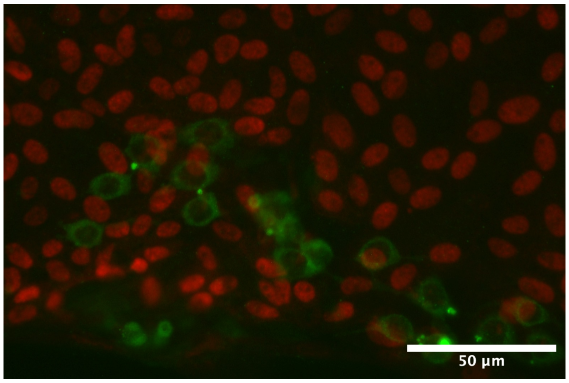
