## Supplemental Figure 6 for "Differential retinoic acid responses across testicular development *in vitro*"

**Supplemental Figure 6. Complete results of transcription factor activity inferenced from bulk RNA-seq data using decoupleR.**

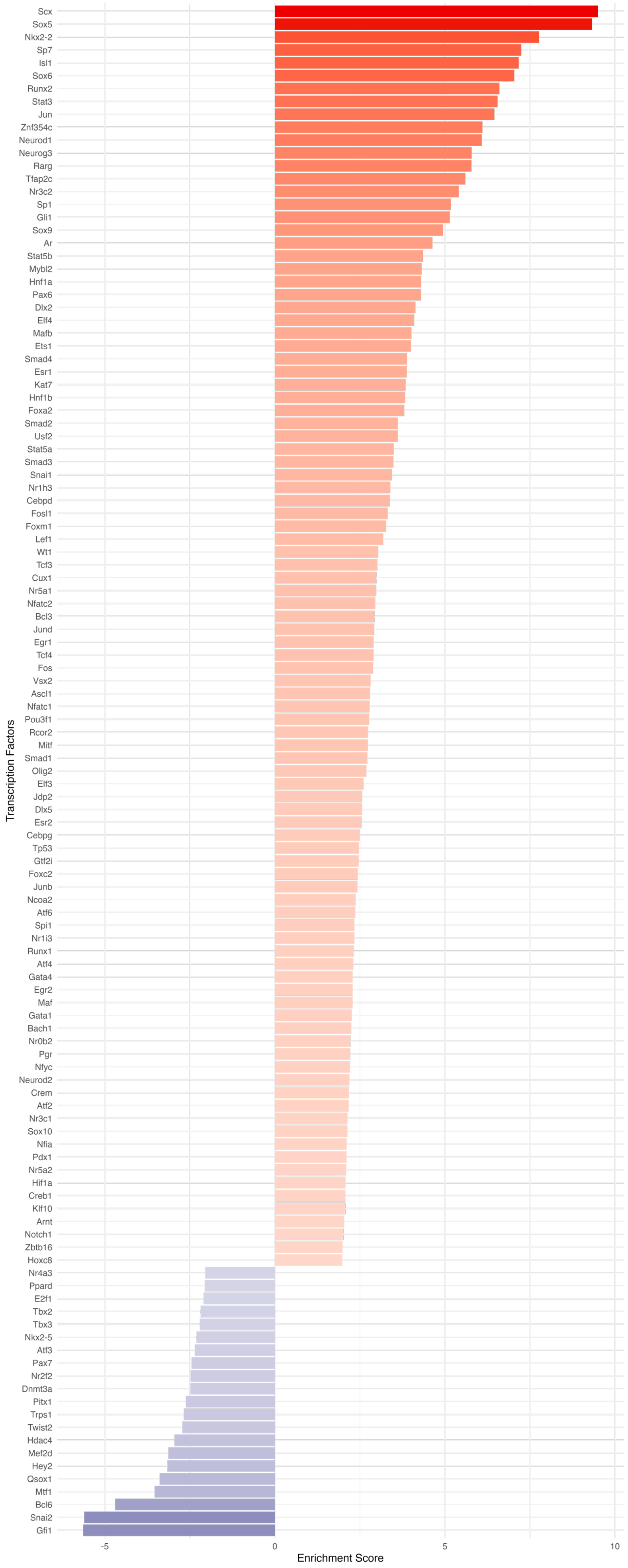
