## Supplemental Figure 8 for "Differential retinoic acid responses across testicular development *in vitro*"

**Supplemental Figure 8. Differentially expressed genes within retinoic acid metabolism related pathways.**

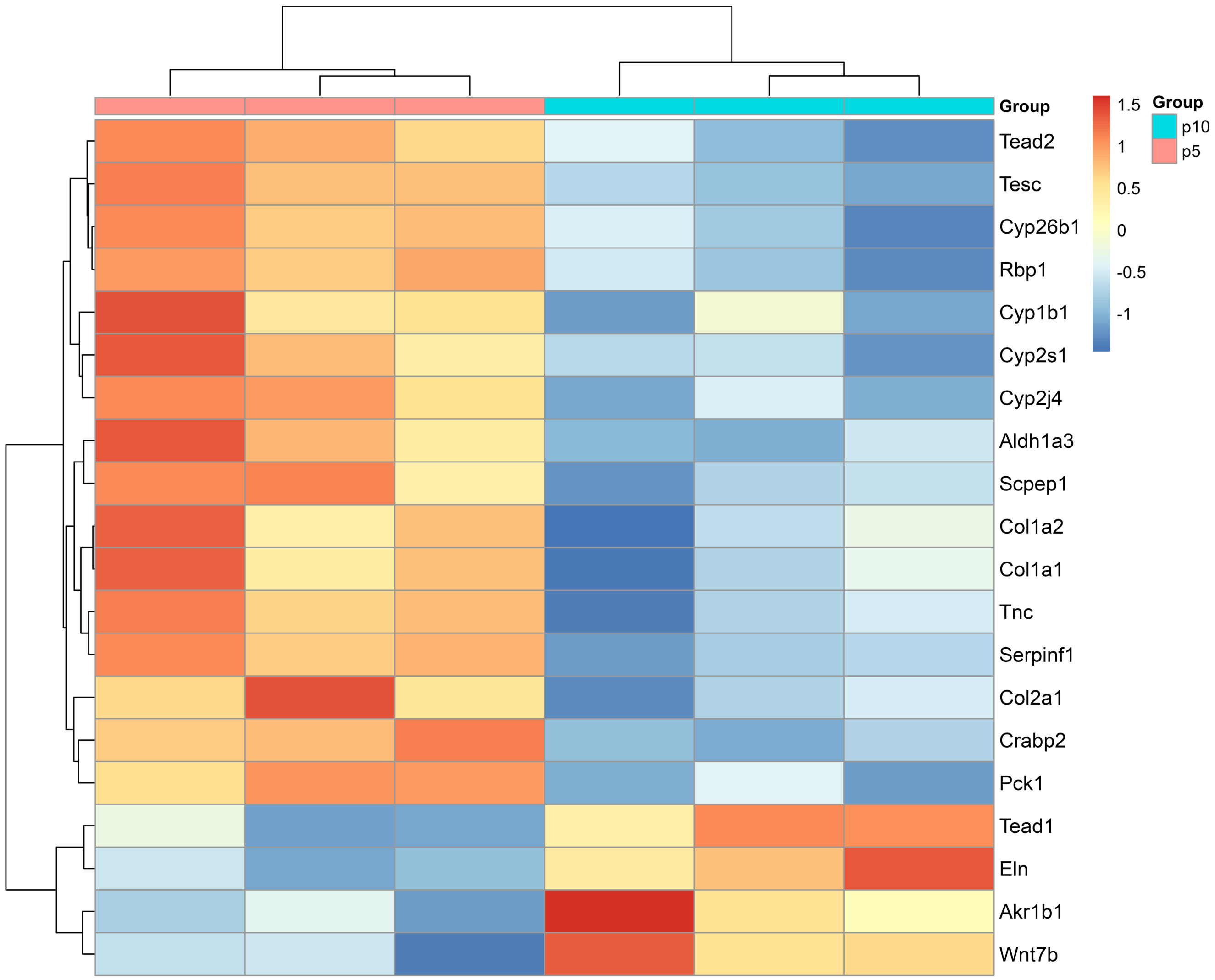
